# Sleep Renormalizes Negative Emotional Generalization

**DOI:** 10.1101/2025.08.18.670780

**Authors:** Ella Bar, Max Bringmann, Gennadiy Belonosov, Kristoffer Aberg, Reut Weissgross, Yuval Nir, Rony Paz

## Abstract

Overgeneralization of negative experiences co-occurs with sleep disruption and characterizes anxiety and posttraumatic stress disorder (PTSD). A strategically-designed behavioral paradigm tested whether sleep shifts emotional generalization away from negative and toward positive representations. Participants learned positive, negative, or neutral face-outcome associations, then completed immediate and delayed generalization testing with morphed faces after sleep or wakefulness. Sleep selectively enhanced positive generalization, whereas wakefulness favored negative generalization. To uncover the underlying neural mechanisms, individual variability in generalization was examined using fMRI and high-density EEG. Amygdala and limbic activity during outcome-processing predicted both stronger immediate negative generalization and a sleep-specific shift toward positive generalization. Additionally, sleep-spindle activity positively correlated with post-sleep increases in positive generalization and reductions in negative generalization. Together, these findings identify a neural mechanism through which sleep reduces negative bias while strengthening positive representations, highlighting a pathway by which sleep may protect against maladaptive generalization associated with anxiety and PTSD.

## Introduction

Imagine walking down the street to your favorite café when a dog suddenly jumps out from around a corner and knocks you to the ground. When you later return to the same street, you encounter another large dog. Even if it only vaguely resembles the original, you may perceive it as the same dog and instinctively retreat. This response reflects emotional generalization—an adaptive process that allows past experiences to guide responses to novel but similar situations^1,2^. Generalization can occur both consciously and unconsciously, shaping how we interpret current stimuli based on prior outcomes ^2,3^. It is especially sensitive to emotionally salient events, with a pronounced bias toward negative associations. For instance, stimuli paired with negative reinforcement tend to produce broader generalization gradients than those associated with positive reinforcement, which in turn generalize more broadly than neutral stimuli^3,4^.

While generalization is essential for adaptive behavior, overgeneralization, and particularly for negative experiences, can become maladaptive. For example, a single dog attack may lead to fear of all dogs or even avoidance of outdoor environments altogether. Such patterns are characteristic of several psychopathologies, including phobias, generalized anxiety disorder, and posttraumatic stress disorder (PTSD)^1,4,5^.

Interestingly, sleep disturbances are also a hallmark of these disorders, and there is a bidirectional relationship between sleep and mental health^6–9^. Disrupted sleep can impair emotional regulation and heighten anxiety, while elevated anxiety can, in turn, degrade sleep quality^10^. Consistent with this, insufficient or disrupted sleep has been shown to bias emotional processing toward negative information, including more negative appraisal of neutral stimuli^11^, impaired disengagement from negative emotional content^12^, heightened reactivity and impulsivity toward negative stimuli^13^, and reduced top-down regulation of emotional responses^14,15^, with broader evidence that sleep loss promotes negatively biased emotional and social processing^9,16^. At the same time, a growing body of research shows that sleep facilitates the consolidation of both rewarding ^17–21^ and aversive^22,23^ emotional memories, while also supporting the extraction of patterns across experiences^24–26^. Given the interplay among sleep, generalization, and mental health, we hypothesized that sleep may serve as a corrective mechanism that regulates emotional generalization by reducing negative bias and restoring a more adaptive balance.

In this study, participants learned associations between neutral faces and positive, negative, or neutral outcomes. Generalization was assessed using a face discrimination task with morphed stimuli, which captured how learned associations extended to novel yet similar inputs. Testing occurred before learning, immediately after learning, and following a consolidation period of either overnight sleep or daytime wakefulness. We found that sleep reduced the generalization of negative stimuli. Results from fMRI and sleep electrophysiology suggest that the amygdala and broader limbic regions, together with spindle oscillations, contribute to sleep-dependent renormalization of generalization, indicating a potential role for sleep in emotional generalization and its relevance for psychopathology.

## Results

To study the role of sleep in the generalization of complex real-life stimuli, we designed a battery of novel behavioral tasks. In a discrimination task, participants (N=60) were presented with face images generated through morphing, depicting intermediate levels along a continuum between three original faces (Figure 1A). For each image, participants had to choose which original face it most closely resembled (Figure 1C). In a Learning task, participants learned to associate the three original faces with either positive, negative, or neutral feedback (Figure 1B, probabilistic two-arm bandit design). All participants performed the discrimination task before the learning task (Block 1) and immediately after it (Block 2). Then, half of the participants were tested on the discrimination task again following an 10-12 hour interval of nighttime sleep, and the other half was tested following a similar interval of wakefulness (Block 3, Figure 1D, Supplementary Figure 2A).

**Figure 1:**
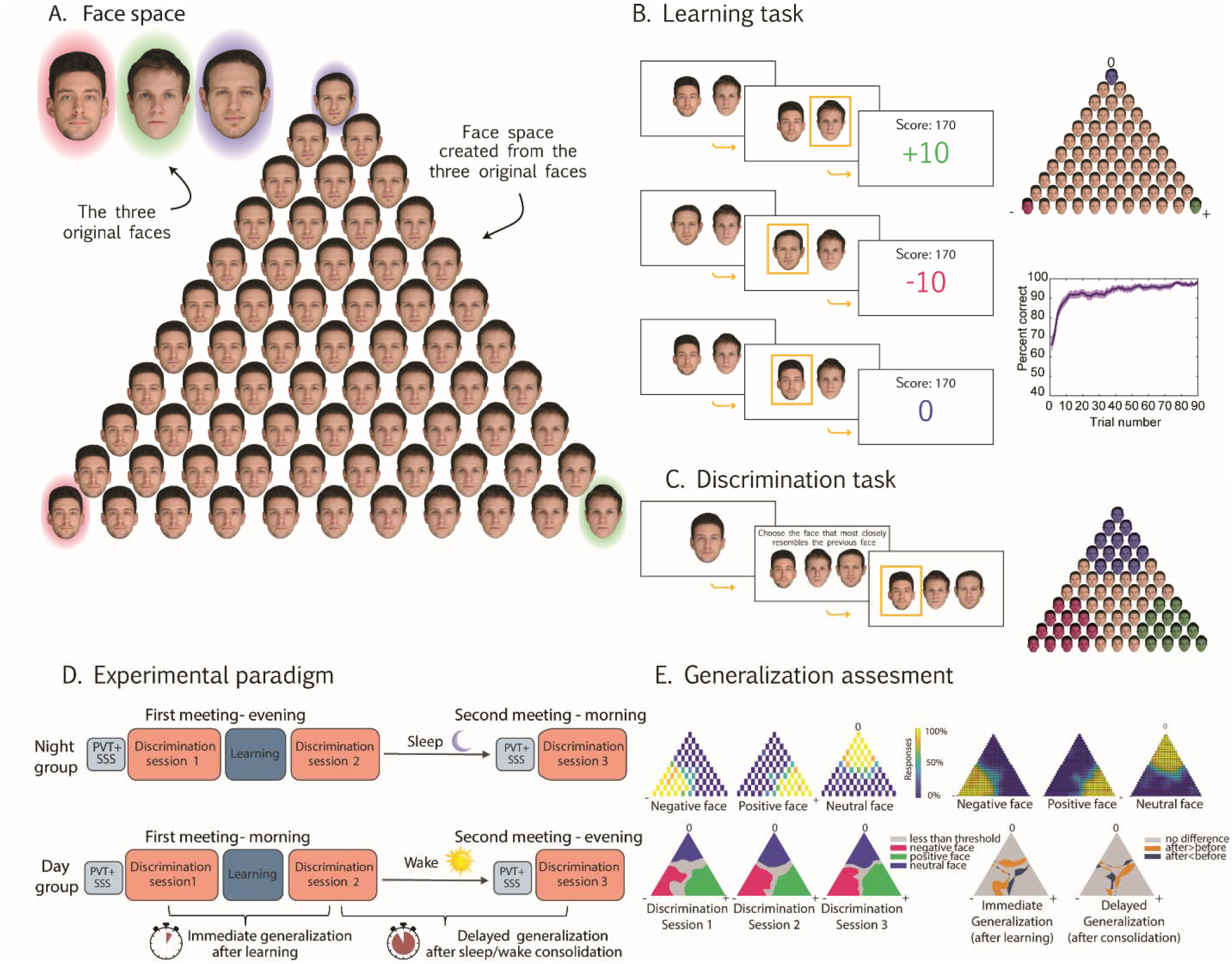
Experimental design: **A.** Face Space: Top left: the three original faces used in the experiment. Right: The triangular "face space" generated from the three original faces. Faces at the vertices are the original faces, while intermediate positions represent morphed faces created by blending the three originals. Each morph’s proximity to a vertex indicates the proportion of that original face in the morph. The face at the center is the average of all three original faces. **B.** Learning task trial timeline. Left: During each trial of the learning task, two faces (selected from the three original faces) were displayed. Participants chose one face and received feedback (+10, -10, or 0 points) based on their selection. Each original face was probabilistically associated with positive, negative, or neutral feedback (90% probability per face for each category). Feedback-face associations were randomized across participants. Detailed timing for different experiments is provided in Supplementary Figure 1. Top-right: Feedback association illustration for the original faces at the vertices of the face space (randomized for each participant). Bottom-right: learning curve from participants across all three experiments (behavioral, fMRI, EEG). **C.** Discrimination task trial. Left: Each trial began with a morphed face from within the face space. Then, participants were asked to choose the original face that most closely resembled the presented morph. Selections were indicated by pressing one of three arrow keys, and the chosen face was highlighted with a yellow rectangle. Detailed timing for different experiments is provided in Supplementary Figure 1. Right: illustration of generalization areas around each original face**. D.** Schematic timeline of the experimental procedure: Participants completed two sessions, approximately 8–12 hours apart, either across the day (Day group) or overnight (Night group). Each session included assessments of sleepiness and vigilance. The first session included two blocks of the discrimination task, separated by the learning task, and the second session included the last block of the discrimination task. In the behavioral and fMRI experiments, participants were assigned to either the Day or Night condition, and sleep or wakefulness between sessions was monitored via actigraphy. In the sleep EEG experiment, only the Night group was tested. (Supplementary Figure 2) **E.** Generalization Assessment Example. Top left: Triangular response space for a single participant, with colors indicating the proportion of responses identifying a morph as similar to the negative (left), positive (middle), or neutral (right) associated face. Top right: Interpolated representation of the triangular response space. Bottom left: Simplified triangular spaces generated by applying thresholds for each original face and merging areas from the same discrimination block into a single triangle. Bottom right: Segment changes across learning – immediate generalization (left). And post-sleep/wake intervals – delayed generalization(right).

Behavioral responses from each discrimination block were organized into a triangular face-space representing perceived similarity between each individual morph and an original face (vertex). To compare generalization across discrimination blocks, we quantified the area around each vertex with higher-similarity judgements (>50% threshold, Figure 1E). In other words, generalization is represented by the overall area of morphed faces classified by a participant as identical to the vertex. Throughout the paper, ’immediate generalization’ refers to differences due to learning and assignment of value to the faces (block 2 - block 1), and ’delayed generalization’ refers to differences due to passage of time, either following sleep or wakefulness (block 3 - block 2).

In a Learning task, participants learned to associate the three original faces with either positive, negative, or neutral feedback (Figure 1B, probabilistic two-arm bandit design). Participants successfully acquired these associations, achieving over 90% accuracy across all experiments and groups (Supplementary Figure 10).

### Sleep promotes generalization of positive associations

We observed differences in generalization patterns following wakefulness and sleep. Following overnight sleep, participants tended to generalize the positive associated face; whereas following wakefulness, participants tended to generalize the negative associated face (Figure 2A). To quantify the effect, we conducted a two-way, mixed-model ANOVA on delayed generalization, with Group (Day/Night, between subjects) and Face (Positive/Negative/Neutral, within-subjects) as factors, and found a significant Face × Group interaction (F (2,116)=3.61, p=0.03, partial η² = 0.059, 90% CI [0.003, 0.132] Figure 2B, 2C). Post-hoc p-values were adjusted for multiple comparisons using the Benjamini–Hochberg procedure across the four follow-up tests. The negative face was generalized more following wakefulness than following overnight sleep (t(58) = -2.3, p = 0.049, d = -0.59, 95% CI [-1.12, -0.07]). In the day group, negative-face generalization was significantly greater than zero (t(29) = 2.44, p = 0.049, dz = 0.44, 95% CI [0.05, 0.84]). In contrast, the positive face was generalized more following sleep than following wakefulness (t(58)=2.01, p=0.049, d = 0.52, 95% CI [-0.01, 1.04]). In the night group, positive-face generalization was also significantly greater than zero (t(29)=2.145, p=0.049, dz = 0.39, 95% CI [0.004, 0.78]). These differences in generalization were not accompanied by group differences in overall discrimination performance. Discrimination accuracy (percentage of correct responses) increased significantly across sessions (χ²(2) = 14.42, p < 0.001), indicating improved task performance over time, but showed no main effect of Group and no Session x Group interaction.

**Figure 2:**
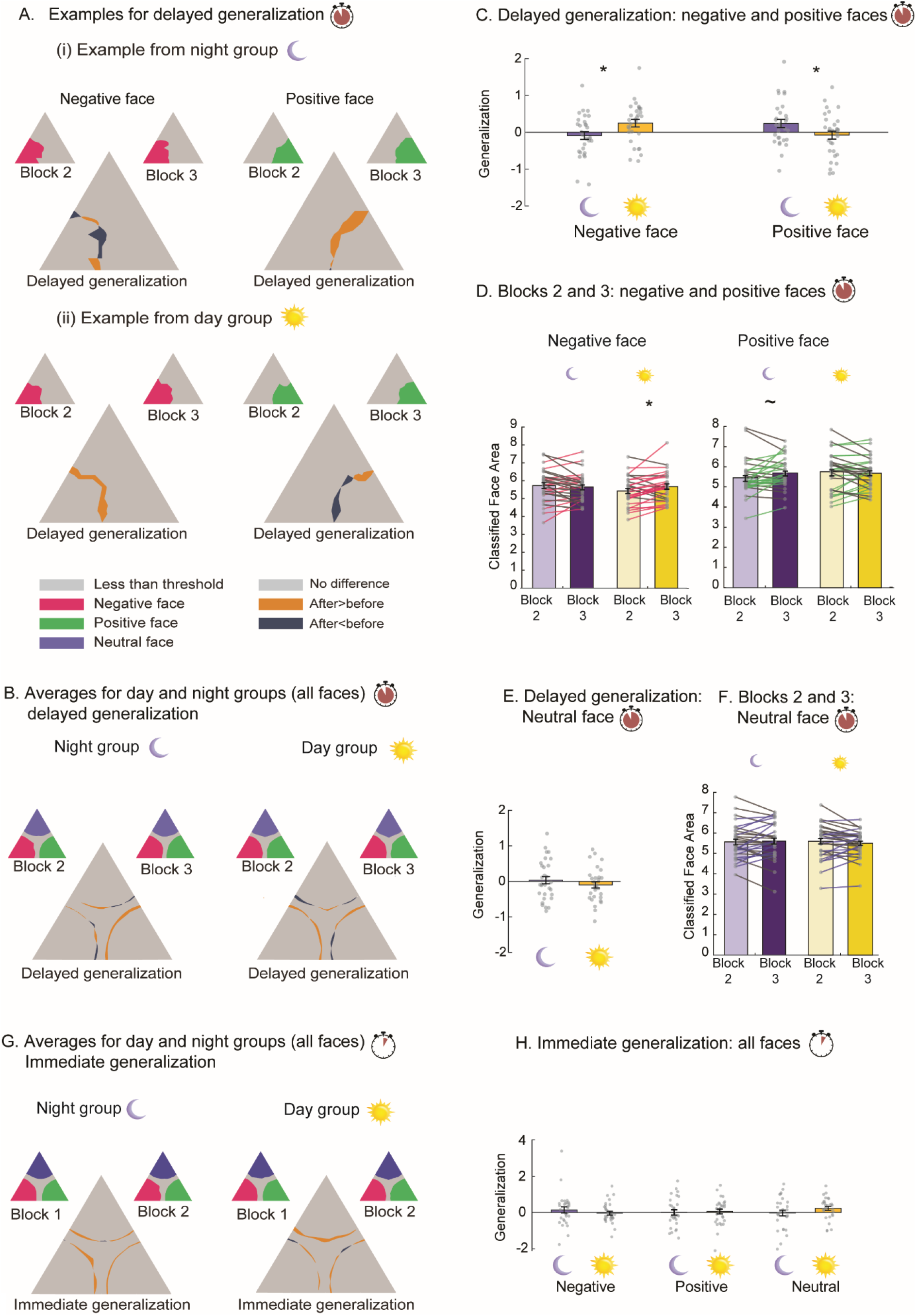
Sleep selectively promotes generalization of positive associations. **A.** Examples of Delayed Generalization from Individual Participants: (i) Night group participant: In the small triangles, pink areas indicate morphed faces classified as the negative face in discrimination blocks 2 and 3; green indicates the positive face. The large triangle illustrates delayed generalization, changes in the generalization areas between blocks 2 and 3 across the night. This participant shows increased generalization of the positive face overnight. (ii) Day group participant: This participant shows delayed generalization of the negative face after a day of wakefulness. **B**. Group-Level Delayed Generalization: Small triangles show averaged morph areas classified as each of the original faces during discrimination blocks 2 and 3, combined onto one triangle. Large triangles show the change in generalization between blocks. Left: night group; right: day group. Night group shows increased delayed generalization of the positive face, day group of the negative face. **C.** Summary of Delayed Generalization: Bar graph shows average change for positive and negative faces from (B). Night group shows significantly greater generalization of the positive face; day group to the negative face. *p<0.05, two-tailed two-sample *t*-test. **D.** Block-wise Change in Classification Area: Bar graph shows size of areas classified as positive and negative in blocks 2 and 3. Day group shows increased negative face classification in block 3. *p<0.05, ∼p<0.10, two-tailed paired *t*-test. **E.** Neutral Face Delayed Generalization: No significant difference between groups. **F.** Block-wise Neutral Face Classification: No significant difference from block 2 to 3 in either group. **G**. Group-Level Immediate Generalization: Same as (B), comparing block 1 and 2. No significant change in any face for either group. **H.** Summary of Immediate Generalization: Bar graph shows average immediate generalization for all faces. No significant differences in either group. Error bars in panels C, D, E, F, and H represent the standard error of the mean (SEM).

To further understand the generalization effect, we conducted separate two-way, mixed-model ANOVAs for the negative and positive faces, this time considering the total classified area in each block (rather than only the block-to-block difference; Figure 2D). For negative face generalization, a mixed-model ANOVA with Group (Day/Night) and Block (Block 2/Block 3) as factors revealed a significant Group × Block interaction (F(1, 58)=5.29, p=0.025, partial η² = 0.084, 90% CI [0.006, 0.214] ; Figure 2D, left). Post-hoc p-values were adjusted for multiple comparisons using the Benjamini–Hochberg procedure. The area classified as negative increased significantly from Block 2 to Block 3 in the daytime wake group (t(29) = −2.44, p = .042, dz = −0.44, 95% CI [−0.84, −0.05]). A similar analysis for positive face generalization also revealed a significant Group × Block interaction (F(1, 58)=4.04, p=0.049, partial η² = .065, 90% CI [0.000, 0.189]; Figure 2D, right). The area classified as positive increased from Block 2 to Block 3 in the sleep group, although this post-hoc comparison did not remain statistically significant after correction for multiple comparisons (t(29) = −2.14, p = .081, dz = −0.39, 95% CI [−0.78, −0.004]). No significant group differences were observed for neutral face generalization (Figure 2E, 2F), nor were there differences in immediate generalization prior to the sleep/wake interval (Figure 2G, 2H). Together, these findings provide preliminary evidence that sleep and wakefulness may be associated with different patterns of emotional generalization, with overnight sleep showing a relative bias toward positive stimuli and daytime wakefulness showing a relative bias toward negative stimuli. However, these effects were modest in magnitude and should be interpreted cautiously.To investigate potential confounds, we compared subjective sleepiness (SSS) and vigilance (PVT) between groups. No significant group differences were found in either measure during the first session. In the second session, participants in the day group reported higher sleepiness than the night group (Day: 3.87 ± 1.63 vs. Night: 3.03 ± 1.34; t(58)=2.15, p=0.036); however, no group differences were observed in vigilance as measured in the PVT (a highly sensitive measure of sleepiness^27,28^). Additionally, neither SSS nor PVT scores were significantly correlated with generalization measures in either group (see Discussion).

### Sleep reverses amygdala-associated generalization of negative stimuli

Next, we set out to investigate the brain networks that contribute to sleep-related generalization of valence-based learning. To do so, an additional group of participants (N=80) performed the learning task while undergoing functional neuroimaging (fMRI, Supplementary Figure 2B). The group-level difference in generalization observed in the behavioral experiment was not replicated in this cohort (Supplementary Figure 3). Discrimination accuracy nevertheless improved across sessions (χ²(2) = 39.63, p < 0.001), with no main effect of Group (and no Session × Group interaction, indicating similar overall discrimination performance across the two groups. There was, however, considerable inter-individual variability in generalization patterns. Such inter-individual variability means that some participants exhibited stronger generalization of the positive face (Figure 3A), whereas others showed a bias toward generalization of the negative face (Figure 3B). This provides a powerful tool for examining brain activity that predicts differences between sleep- and wakefulness-based generalization.

**Figure 3.**
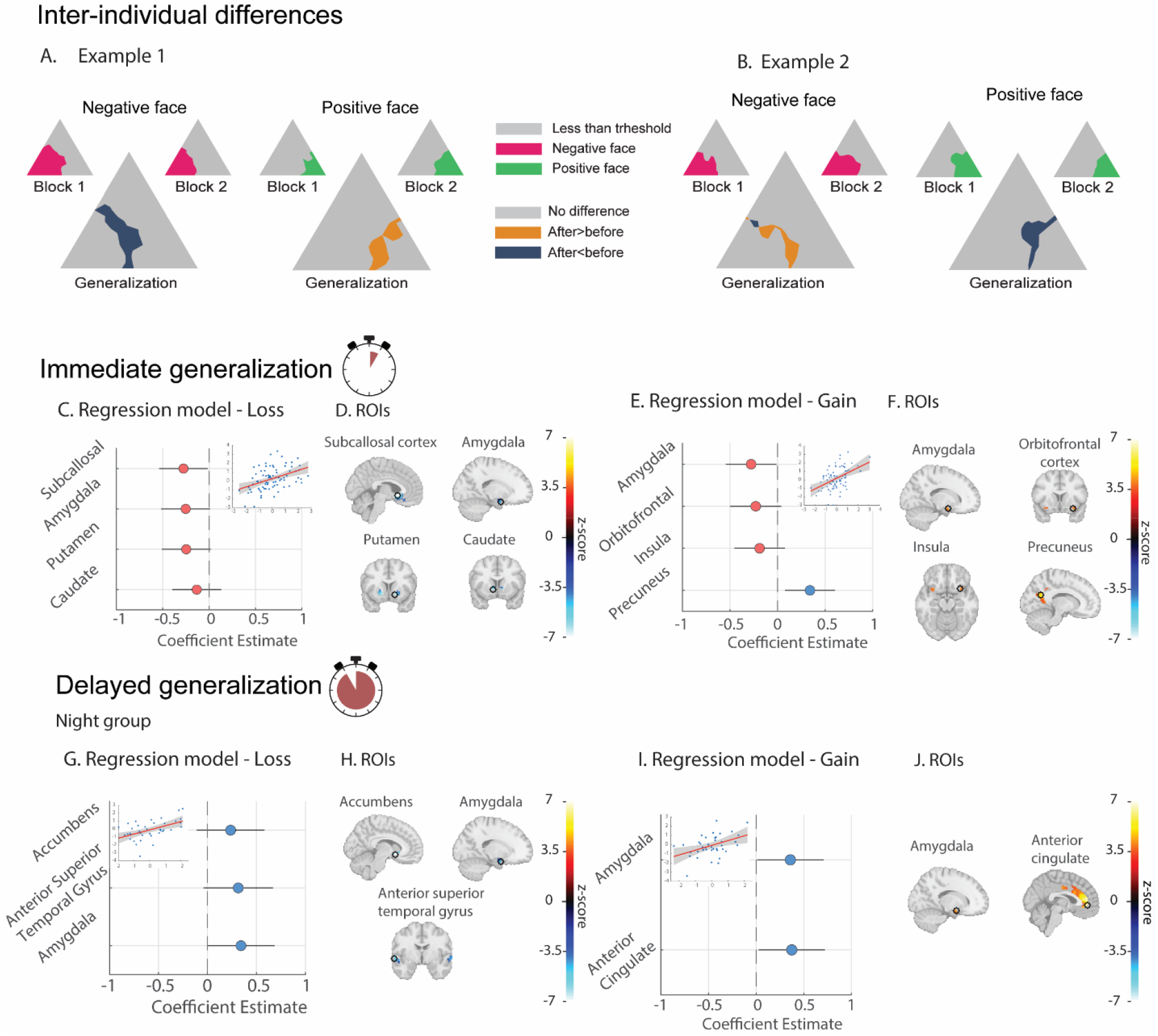
**Immediate and delayed generalization are correlated with the amygdala and limbic activity. A–B**. Individual variability in immediate generalization. **A.** A participant who generalized the positive face immediately after learning. Left: areas classified as the negative face during discrimination blocks 1 and 2. Right: areas classified as the positive face. Larger triangles represent changes in classification between blocks, reflecting immediate generalization. **B.** regions classified as the negative face during blocks 1 and 2. Right: regions classified as the positive face. **C-D.** Regression results for predicting immediate generalization from neural activity during loss-related feedback. **C.** Coefficient estimates for ROIs selected via Lasso regression: Subcallosal Cortex, Amygdala, Putamen, and Caudate. All coefficients are negative, indicating that greater activity in these limbic regions was associated with reduced generalization of the positive face and increased generalization of the negative face. Inset (top right): model fit (x-axis=predicted generalization from ROI activity; y-axis=observed generalization). Red line=regression fit; gray shading=95% confidence intervals. **D**. Mean ROI activity during loss feedback for regions in (C). Black circle visualize a 3-voxel radius sphere centered on the peak voxel within each anatomical ROI (for visualization only). **E-F**. Immediate generalization: gain-related feedback. **E.** ROIs selected via Lasso included the amygdala, orbitofrontal cortex, insula, and precuneus. Limbic regions showed negative coefficients, indicating that greater activity was associated with reduced positive-face generalization. Precuneus (DMN) activity was positively associated with the generalization of the positive face. Inset as in C. **F.** Mean ROI activity during gain feedback for regions in (E). **G-H.** Delayed generalization in the night group: loss-related feedback. **G.** Coefficient estimates for ROIs selected via Lasso regression: Accumbens, Anterior Superior Temporal Gyrus, and Amygdala. All coefficients are positive, indicating that increased activity in these regions was associated with greater generalization of the positive face and less generalization of the negative face. Inset (top left) as in C. **H.** Mean ROI activity during loss feedback for regions in (G). I-J. Delayed generalization: gain-related feedback. **I.** The amygdala and anterior cingulate cortex were selected via LASSO. Both coefficients are positive, suggesting that increased activity in these regions was associated with greater positive-face generalization and decreased negative-face generalization. Inset (top left) as in C. **J.** ROI activity during *gain* trials for regions shown in panel I.

First, we tested whether brain activity during learning, specifically when receiving feedback linked to point gains or losses, could predict immediate and delayed generalization behavior. To quantify generalization, we computed a composite score calculated as the difference between positive- and negative-face generalization scores (Methods), with values above zero indicating greater generalization of the positive face and below zero indicating greater generalization of the negative face. This composite measure was selected as the primary outcome because brain activity showed opposing associations with positive- and negative-face generalization, and the difference score therefore provides a parsimonious summary of the direction and magnitude of emotional generalization bias. This score was used in multiple linear regression models to predict either (i) immediate generalization (for both groups combined, as it is prior to the sleep/wake consolidation period) or (ii) delayed generalization following daytime wake or overnight sleep (separated between the groups). Although the composite score served as the primary outcome measure for simplicity of presentation, separate models predicting positive- and negative-face generalization independently yielded the same qualitative pattern of results and are provided in Supplementary Figures 4 and 5. fMRI blood oxygenation level dependent (BOLD) activity underwent whole-brain general linear model (GLM) analysis to identify task-related activity, and 14 Regions of Interest (ROIs) were selected as predictors (Methods). To identify the most predictive ROIs, we applied Lasso regression ^29^ (elastic net regularization) with 10-fold cross-validation for model optimization.

**Figure 4.**
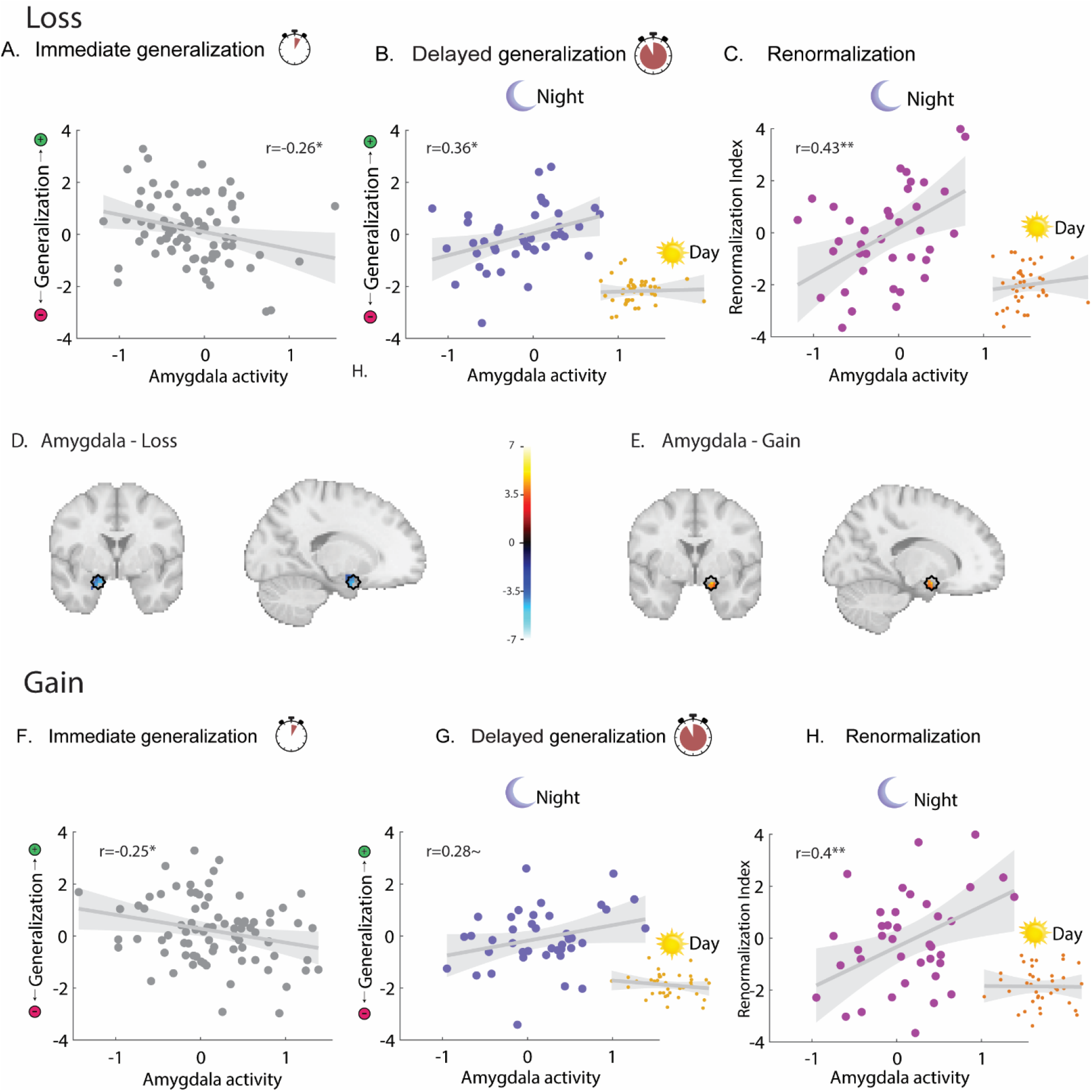
Direct correlations between amygdala activity during learning and generalization behavior. **A.** Immediate generalization: Amygdala activity during loss feedback in the learning phase is negatively correlated with immediate generalization across both groups. Higher amygdala activity is associated with increased generalization of the negative face. The grey line represents the fitted regression line; the gray shaded area indicates 95% confidence intervals. **B.** Delayed generalization. Left: In the night group, higher amygdala activity during loss is associated with positive face generalization after sleep. Bottom right: No such correlation is observed in the day group, suggesting the effect is sleep-specific. **C.** Correlation with the renormalization index. Left: In the night group, higher amygdala activity during loss is significantly correlated with a greater renormalization index, indicating a stronger shift from negative to positive generalization after sleep. Bottom right: No such relationship is observed in the day group. **D.** Activation map showing the amygdala response to loss feedback. The black circle visualize a sphere (radius=3 mm) centered on the peak activation voxel (size is for visualizasion only). Activity used for analysis is the average across voxels within this sphere that fall within the anatomical ROI. **E.** Same as D for gain feedback. **F–H**. Same analyses as in panels A–C, but for amygdala activity during gain feedback. **F**. Higher amygdala activity during gains is associated with increased immediate generalization of the negative face. **G.** Left: Amygdala activity during gains predicts a renormalization toward positive-face generalization after sleep in the night group. Bottom right: No such effect is observed in the day group. **H.** Correlation between amygdala activity during gains and the renormalization index: significant in the night group (left), absent in the day group (bottom right). **p<0.01, *p<0.05, ∼p<0.10, Pearson’s r.

**Figure 5:**
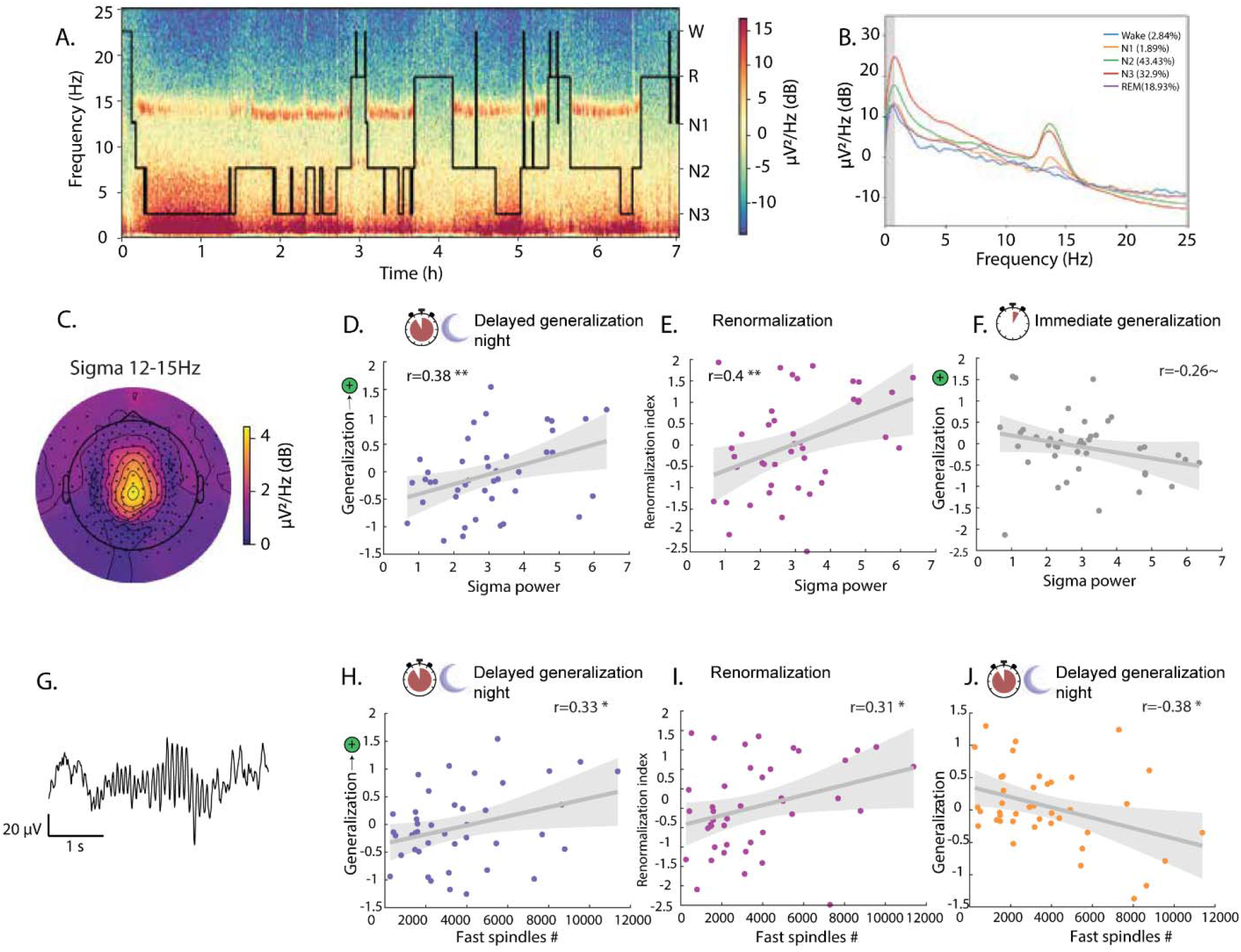
**Sleep spindle activity facilitates positive generalization**. **A.** Time–frequency spectrogram (log power) from a representative participant recorded at electrode Pz, illustrating spectral dynamics across the night. The overlaid hypnogram depicts transitions between sleep stages. **B.** Power Spectral Density (PSD) plots from preprocessed data at electrode Pz from a representative participant, separated by sleep stage. Colored lines denote individual stages. The gray-shaded region indicates frequencies removed by the high-pass filter. **C.** Topographical map of sigma power (12–15 Hz) during N2 sleep from a representative participant, demonstrating the typical centroparietal distribution of sleep spindles. **D.** Sigma power during NREM sleep is positively correlated with delayed generalization of the positive face measured after sleep. **E.** Sigma power during NREM sleep is significantly correlated with the renormalization of positive face generalization (i.e., the change from pre-sleep/immediate to post-sleep/delayed generalization). **F.** Correlation between sigma power during NREM sleep and immediate generalization of the positive face. Participants with lower immediate positive generalization exhibited greater subsequent sigma power. **G.** Representative example of a fast spindle (13–15 Hz) recorded during NREM sleep in the time domain. Scale bars indicate 1s (horizontal) and 20µV (vertical). **H.** The number of fast spindles during NREM sleep is positively correlated with delayed generalization of the positive face. **I.** Fast spindle count is positively correlated with the renormalization of positive generalization from immediate to delayed generalization. **J.** Greater spindle count during NREM sleep is associated with reduced delayed generalization of neutral faces, suggesting selective suppression of non-valenced stimulus generalization. **p<0.01, *p<0.05, ∼p<0.10, Pearson’s r.

We found that greater activation for losses (versus neutral feedback) in limbic regions, specifically the Amygdala, Caudate, Putamen, and Subcallosal Cortex, was correlated with increased immediate generalization of the negative face, but decreased immediate generalization of the positive face, This corresponds to a negative correlation with the generalization composite score (regression model: N=80, R²=0.18, F(4,75)=4.17, p=0.004; Figure 3C, D). A similar pattern emerged during gains (versus neutral feedback): activity in the Amygdala, Orbitofrontal cortex, and Insula was positively correlated with generalization of the negative face and negatively with the positive face (positive correlation to the generalization score), whereas activity in the Precuneus (default mode network, DMN), showed the opposite trend: (regression model: N=80, R^2^=0.18, F(4,75)=4.14, p=0.004, Figure 3E, F). Thus, regardless of feedback valence (gaining or losing points associated with monetary reward), limbic activity biases immediate generalization toward negatively associated stimuli, whereas engagement of the DMN is associated with positive stimuli’s immediate generalization. These effects were also evident when modeling positive and negative generalization separately, rather than using the generalization composite score. Specifically, limbic activity during learning mostly predicted increased immediate generalization of the negative face (see Supplementary Figure 4).

Next, we analyzed brain activity associated with delayed generalization after overnight sleep. Greater limbic activation during both loss- and gain-related feedback in the learning task was associated with a renormalization from negative to positive face generalization following overnight sleep. Specifically, activity in the Amygdala, Nucleus Accumbens, and Anterior Superior Temporal Gyrus during loss-related feedback predicted increased generalization of the positive face after sleep (regression model: N=40, R²=0.27, F(3,36)=4.45, p=0.009; Figure 3G, H). A similar trend was observed for gain-related feedback, where higher activation in the Amygdala and Anterior Cingulate Cortex during gains was associated with a greater shift in generalization from the negative to the positive face following sleep (regression model: N= 40, R²=0.18, F(2,37)=4.14, p=0.02; Figure 3, I-J). Similarly, separate models for delayed positive and negative generalization, instead of the generalization composite score, showed that limbic activation during learning predicted a sleep-dependent shift toward greater generalization of the positive face (see Supplementary Figure 5).

Together, these findings suggest that heightened limbic engagement during emotionally salient learning, regardless of outcome valence, may promote immediate negative generalization, followed by sleep-dependent renormalization that shifts toward positive generalization. In contrast, applying the same Lasso regression approach to predict delayed generalization after daytime wakefulness did not select any ROI predictors for either gain or loss, suggesting that this renormalization process may be specific to sleep-related consolidation (see Discussion).

Importantly, the pattern of results remained unchanged after controlling for subjective sleepiness (SSS) and vigilance (PVT). The group effect (χ²(1)=5.81, p=.016) and the face-by-group interaction (χ²(2)=7.67, p=.022) remained significant, whereas neither SSS (χ²(1)=2.13, p=.144) nor PVT (χ²(1)=0.61, p=.433) significantly predicted generalization.

Amygdala activity consistently emerged as an important predictor of immediate and delayed generalization when activated for both positive and negative emotional valence stimuli, being the only ROI repeatedly selected by LASSO across all prediction models. Specifically, amygdala activation during losses was significantly correlated with immediate generalization of the negative face (negative correlation to the composite score) across both experimental groups (N=80, r_p_=-0.26, p=0.02, Figure 4A). However, for delayed generalization following overnight sleep, this activity predicted the positive face generalization (N=40, r =0.36, *p*=0.02; Figure 4B), a phenomenon that did not occur after daytime wakefulness, although Fisher’s r-to-z comparisons between groups were not significant. (Figure 4B, bottom-right, r_p_=0.04, p=0.8, Fisher test to compare night and day correlations: Z=1.45, p=0.15).

To quantify this effect, we computed a renormalization index for each participant by subtracting the immediate generalization score from the delayed generalization score for each participant (Methods). Positive renormalization indices indicate that, after sleep, generalization shifts towards more positive values than at baseline. In the night group, greater amygdala activation during loss-related feedback predicted stronger renormalization following sleep. Participants with higher amygdala activity not only exhibited greater immediate generalization of the negative face but also demonstrated a clear shift toward positive face generalization after sleep, as reflected by a significant correlation between the renormalization index and amygdala activity during loss (N=40, r =0.43, *p*=0.005; Figure 4C). No such correlation was observed in the daytime group, direct comparison between groups was not significant. (Figure 4C, bottom-right, r =0.12, p=0.45, Fisher test to compare night and day correlations: Z=1.46, p=0.14).

A similar pattern emerged when examining amygdala activity during reward-related feedback. Higher amygdala activation while gaining points was associated with greater immediate generalization of the negative face across both groups (N=80, r =-0.25, p=0.02; Figure 4F). Following sleep, amygdala activity was positively correlated with positive-face generalization, although this association did not reach statistical significance (N = 40, r = 0.28, p = 0.08; Figure 4G). No corresponding association was observed in the daytime wake group (r = −0.16, p = 0.31), and Fisher’s r-to-z comparison indicated a significant difference in correlation strength between groups (Z = 1.90, p = 0.03; Figure 4G, bottom-right).

Furthermore, the renormalization index showed a strong correlation with Amygdala activity during gains (N=40, r =0.4, p=0.009; Figure 4H), indicating that individuals with higher initial amygdala engagement for gain-related feedback also underwent a greater shift from negative to positive generalization after sleep. As observed for loss-related activity, renormalization effects were specific to overnight sleep; there was no correlation with the renormalization index in the daytime wake group (4H, bottom, right panel; r =-0.01, p=0.95, Fisher test to compare night and day correlations: Z=1.87, p=0.06).

Overall, these findings suggest that amygdala activity during learning is associated with subsequent sleep-dependent shifts in emotional generalization, although group differences in correlation strength were not uniformly significant across analyses.

Sleep-dependent renormalization was consistently observed for Amygdala activity, but not unique to this brain region. Direct correlations with other limbic ROIs revealed a consistent pattern in which higher activation during learning (either reward or loss) was associated with greater immediate generalization of the negative face, followed by renormalization toward the positive face after sleep (Supplementary Figures 6 and 7).

In addition, we examined overgeneralization, defined as generalization that is no longer adaptive (i.e., leads to point loss). We found significant correlations between amygdala activity during both loss- and gain-related feedback and two behavioral outcomes: immediate overgeneralization of the negative face and delayed overgeneralization of the positive face following sleep. Notably, overgeneralization scores were highly correlated with overall generalization scores, and the observed associations with amygdala activity did not remain significant after controlling for this shared variance (Supplementary Figure 8).

### Sleep spindle activity correlates with renormalization of negative-to-positive emotional generalization

To further investigate the role of sleep in promoting positive generalization and reversing negative emotional bias, we conducted an overnight polysomnography sleep study with high-density (256-channel) EEG in a separate cohort of healthy volunteers (n=42, Methods, Figure 5A, B). Participants completed the same behavioral paradigm used in prior experiments, both before and after sleep (Supplementary Figure 2C). Discrimination accuracy increased significantly across sessions (χ²(2) = 28.24, p < 0.001), indicating improved task performance over time, no main effect of Group and no Group x Session interaction. Sleep scoring was performed to assess whether sleep architecture was related to generalization outcomes. On average, participants exhibited a typical profile for overnight sleep in healthy young adults, with a mean (SD) total sleep time of 400.2 (69.7) min, sleep efficiency (sleep time/time in bed) of 92.7% (5.2), and a mean sleep latency of 18.3 (15.5) min. Non-rapid eye movement (NREM) sleep, REM sleep, and wake after sleep onset (WASO) constituted 302.2 (32.3), 96.8 (32.3), and 19.7 (17.7) minutes, respectively. We tested for associations between time spent in specific sleep stages and generalization, but no significant relationships emerged (Supplementary Table 1). Given previous reports linking REM fragmentation to impaired emotional recalibration, we additionally examined REM continuity and transient high-frequency arousals during REM sleep. Neither measure was significantly associated with overnight generalization outcomes (Supplementary Table 1). Next, we examined whether specific electrophysiological activities during sleep were associated with delayed generalization. We quantified spectral power in the delta (0.75–4 Hz) and sigma (12–15 Hz) bands during NREM sleep, and theta power (4–8 Hz) during REM sleep (Methods; Supplementary Table 1, Supplementary Figure 9). While delta and theta power did not significantly correlate with delayed generalization measures, sigma-band activity during NREM sleep showed the strongest association with positive-face generalization and was also generally consistent with the broader renormalization pattern across measures. We therefore focused the main text on positive-face generalization as the primary sleep-related effect. For completeness, Supplementary Table 1 reports the associations between all sleep parameters and the generalization of each face type individually, as well as the combined generalization bias score.^30^ Sigma band that is associated with sleep spindles have been previously shown to play a central role in memory consolidation and cortical plasticity^31–35^ . We found that greater sigma power during NREM sleep was significantly correlated with increased delayed generalization of the positive face following sleep (r_p_*=*0.38, p=0.01; Figure 5D). Consistent with this effect, the renormalization of positive face generalization (i.e., the change from immediate to delayed testing) was also significantly correlated with sigma power during sleep (r_p_=0.40, p=0.009; Figure 5E).

The association with the renormalization index suggests that sleep spindles may be related not only to positive generalization itself, but also to changes in emotional generalization across sleep. One possibility is that spindle activity is involved in the rebalancing of emotional generalization when a negative bias is present before sleep. Consistent with this possibility, individuals who initially generalized the negative face more strongly, and the positive face less strongly, showed numerically higher sigma power during subsequent sleep (negative face: r_p_=0.25, p=0.09; positive face: r_p_=−0.26, p=0.09; Figure 5F). However, these associations did not reach statistical significance and should therefore be interpreted cautiously.

To extend this analysis, we went beyond power spectral analysis to detect and quantify individual spindle events using SleepEEGpy^36^, incorporating YASA^37^ detection (Methods, Figure 5G). We found that the number of fast (13–15 Hz) spindles during NREM sleep was positively correlated with post-sleep generalization of the positive face (r =0.33, p=0.03; Figure 5H) and with the renormalization in positive generalization (r =0.031, p=0.04; Figure 5I). Notably, we also found that a higher number of fast spindles was associated with reduced generalization of the neutral face after sleep (r =–0.38, p=0.013; figure 5J). Since neutral faces were not associated with reward or punishment, this finding suggests that sleep spindles may help suppress the generalization of emotionally irrelevant or neutral stimuli, highlighting a possible role of spindles in valence-selective memory processing (see Discussion).

## Discussion

In this study, we tested the hypothesis that sleep reduces emotional generalization of negative stimuli while boosting the generalization of positive stimuli. Using a face discrimination task with morphed faces, we found that in the behavioral experiment overnight sleep selectively promoted generalization of positive associations, whereas daytime wakefulness was linked to generalization of negative associations (Figure 2). While this group-level pattern was not replicated in the fMRI experiment, analysis of fMRI data nevertheless revealed that greater limbic responses to emotional feedback during learning (gains or losses) were associated with broader immediate generalization of the negative face. In the sleep group, the strength of this activity was also associated with a reversal of this pattern after sleep: participants exhibited greater generalization of positive faces following overnight sleep, but not after a day awake (Figure 3). Amygdala activity emerged as an important predictor in several of these analyses, including immediate negative generalization and sleep-associated changes in generalization (Figure 4). Participants with high amygdala activation during learning exhibited the strongest immediate generalization of the negative face and greater renormalization across sleep. Finally, we found that sleep spindle activity was significantly correlated with a shift from negative to positive generalization (Figure 5). Taken together, these findings are consistent with a role for sleep in the reorganization of emotional generalization, and also indicate that the strength and sleep-specificity of these effects vary across measures and experiments.

Data from the behavioral experiment revealed a clear divergence in delayed generalization profiles following overnight sleep or daytime wakefulness. Participants who remained awake after learning tended to generalize the negatively associated face, reinforcing a negative bias, while those who slept were more likely to generalize the positively associated face. These findings align with previous research showing that negative stimuli often elicit broader generalization curves than neutral or positive stimuli ^1,3,4^, a pattern thought to reflect an adaptive threat-detection mechanism. Our results extend this literature by showing that sleep does not merely preserve these biases but actively reshapes the generalization landscape, attenuating negative overgeneralization and enhancing positive generalization

fMRI data showed that greater amygdala activity during processing of either positive or negative outcomes was linked to broader generalization of negative stimuli immediately after learning. After sleep, this pattern reversed: higher amygdala activity was now associated with greater generalization of positive faces, suggesting that sleep supports a renormalization of amygdala-related emotional memory. This effect was absent in the wake group, reinforcing its sleep dependency. Notably, participants with high amygdala activation showed, after sleep, even greater positive generalization than at baseline, extending beyond simple renormalization. These findings align with previous research linking amygdala activation to loss encoding and the broadening of generalization curves for negatively associated stimuli^1,4^. Moreover, sleep deprivation has been shown to amplify amygdala reactivity to emotional stimuli and impair discrimination between threat and safety cues ^9,38^, potentially contributing to overgeneralization in anxiety disorders. Our results suggest that sleep may help restore the negative bias by shifting amygdala-linked generalization away from negative and toward positive stimuli.

To investigate the role of sleep-specific neurophysiology in this reorganization, we recorded high-density EEG during overnight sleep. We found that both sigma power and the number of fast spindle events during NREM sleep were positively correlated with the generalization of positively associated faces. Both measures of spindle activity were significantly correlated with the renormalization in positive face generalization, reflecting a shift from negative to positive bias after sleep. These findings extend our understanding of the association between sleep spindles and memory consolidation, showing that spindles play a role in the emotionally adaptive reorganization of generalized learned associations. In addition to modulating emotional memory, our results suggest that sleep spindles may contribute to the selective suppression of emotionally neutral information. Specifically, greater fast spindle activity was associated with reduced generalization of neutral faces, which were not linked to reward or punishment. This pattern suggests that spindles may serve a dual function, supporting the generalization of emotionally salient memories while promoting forgetting of behaviorally irrelevant content. This interpretation extends prior work proposing that sleep supports the prioritization of salient or goal-relevant information during consolidation^18,22,39^.

One possible neurophysiological mechanism underlying these effects relates to the inverse relationship between NREM spindle activity and arousal systems. During NREM sleep, spindle activity is anti-correlated with markers of arousal, including locus coeruleus–norepinephrine (LC–NE) activity, corticotropin-releasing hormone (CRH) signaling, heightened autonomic tone, and increased microarousals^40–43^. Thus, beyond their role in memory consolidation, increased spindle activity may also index a state of reduced noradrenergic tone. Such a “silent” noradrenergic signaling may provide a permissive environment for the renormalization of negatively biased generalization, allowing previously heightened negative representations to be renormalized toward a more adaptive balance. Within this framework, spindles may not only gate hippocampo–cortical communication, but also reflect a broader physiological state that facilitates emotional recalibration during sleep.

These findings carry important translational implications. While both negative overgeneralization^1,4,50–52^ and disrupted sleep^6,8,9,53–55^ have been linked to anxiety and PTSD, few studies have examined how they interact to shape emotional memory and vulnerability to psychopathology ^23^. Our results suggest that sleep, particularly NREM spindle activity, may serve as a protective mechanism. Sleep spindles may not only reduce negative amygdala-related overgeneralization but also promote adaptive generalization of positive content. Previous studies have shown that sleep spindle density increases following stress^56^ and is elevated in individuals with PTSD^57–59^. While trauma may trigger these increases, our data suggests that spindles may also play a role in emotional recovery. Indeed, sleep spindles in individuals with PTSD have been linked to healthier sleep, reduced anxiety, and improved emotional recovery^56,60–62^. In contrast, lower spindle density has been associated with greater vulnerability to sleep disturbances following stress^67^. Future work could build on these findings, testing whether enhancing sleep, and particularly sleep spindles, can reduce emotional overgeneralization in clinical populations. Promising interventions could include targeted memory reactivation (TMR), which reintroduces learning cues during sleep to reactivate specific memories^35,63–65^, brain stimulation techniques designed to boost sleep spindle activity^66,67^, and pharmacological treatments to elevate sleep spindle activity^68,69^. Applying these methods to emotional generalization paradigms could help advance sleep-based interventions for anxiety and PTSD.

### Limitations of study

Some limitations of this study should be explicitly acknowledged. First, circadian factors may influence the observed group differences. Because testing sessions occurred at opposite times of day (morning vs. evening), it is difficult to fully disentangle the effects of sleep from those of time of day. In the behavioral experiment, participants in the day group reported higher subjective sleepiness in the evening session than the night group did in the morning, although objective vigilance (PVT) scores did not differ. Previous research has linked fatigue to more negative interpretation biases^9,38^, which may in principle partly explain the negative generalization in the day group. However, neither SSS nor PVT scores significantly predicted generalization, and the main effects remained significant after statistically controlling for both measures. Similarly, no group differences in either subjective sleepiness or vigilance were observed in the fMRI experiment. Further arguing against circadian confounds, robust correlations between individual sleep spindle activity in the sleep EEG experiment and shifts in emotional generalization support the interpretation that sleep plays an active, corrective role in modulating emotional overgeneralization. Such renormalization may be mediated in part via spindle-related mechanisms operating during NREM sleep.

Second, group-level generalization differences in the behavioral experiment were not replicated in the fMRI experiment, where no group differences in subjective sleepiness were found. While we can only speculate about the source of this discrepancy, the two paradigms differed substantially in both learning duration and temporal structure. The fMRI experiment involved over one hour of learning, compared with approximately 15 minutes in the behavioral task, and included substantially longer intervals between choices and feedback to allow separation of neural responses. Delayed feedback has been shown to alter learning dynamics and subsequent memory consolidation, even when the reinforcement itself remains unchanged^44,45^. Consistent with this possibility, participants exhibiting stronger amygdala responses during learning in the fMRI experiment showed stronger immediate negative generalization and stronger sleep-dependent shifts toward positive generalization, qualitatively resembling the group-level pattern observed in the behavioral experiment. Thus, procedural differences in the fMRI task may have weakened the average behavioral effect while preserving substantial individual differences linked to amygdala engagement^44^.

Third, the reinforcement outcomes used in the present study were substantially less aversive than the traumatic experiences that characterize anxiety- and stress-related disorders. Previous work has demonstrated that outcome intensity can influence generalization processes, with stronger aversive outcomes producing broader and less selective generalization gradients^46^. Although threat intensity was not manipulated in the current study, participants with stronger amygdala responses during learning exhibited greater immediate negative generalization, suggesting that individual differences in neural salience may play a role analogous to outcome intensity. This interpretation is consistent with previous work implicating the amygdala in emotional salience and fear generalization^1,47^. Interestingly, these same individuals also showed the strongest sleep-dependent shift toward positive generalization. This raises the possibility that sleep may preferentially renormalize highly salient emotional memories, consistent with theories proposing that sleep selectively processes emotionally significant experiences^48,49^. Future studies should examine whether similar sleep-dependent renormalization processes occur following more intense aversive experiences and in clinical populations characterized by maladaptive overgeneralization.

An additional limitation is that sample sizes were determined primarily based on feasibility considerations and prior literature benchmarking rather than pilot-based power analyses. Sensitivity analyses indicated that the present studies were most sensitive to effects in the moderate range, and therefore smaller effects may not have been reliably detectable. Accordingly, null findings should be interpreted with appropriate caution, particularly for analyses where the true effects may be small.

## STAR Methods

### Resource availability

#### Lead contact

Further information and requests for resources should be directed to and will be fulfilled by the lead contacts: Prof. Rony Paz or Prof. Yuval Nir.

## Materials availability

This study did not generate new unique reagents or materials.

## Data and code availability

### Data

The data supporting the findings reported in this study have been deposited in Mendeley Data and are publicly available at 10.17632/s9vxcsk9yx.1.

### Code

The original code used for the analyses reported in this study has been deposited on GitHub and is publicly available at https://github.com/elba1212/Sleep_Generalization_Code.

### Additional information

Any additional information required to reanalyze the data reported in this paper is available from the lead contact upon request.

## Experimental model and study participant details

### Behavioral Experiment

#### Participants

Sixty healthy participants took part in the behavioral experiment, randomly assigned to a day group (n=30) or a night group (n=30). Written informed consent was obtained from each participant, and the study was approved by the IRB of the Weizmann Institute of Science. No participants were excluded. The final sample included 30 participants in the night group (19 females, mean age=25.57, SD=3.11) and 30 in the day group (14 females, mean age=28.13, SD=6.4). No pilot dataset was available. The target sample size was determined prior to data collection based on feasibility and resource constraints, benchmarking prior literature, and the goal of exceeding sample sizes commonly used in related studies of sleep-dependent emotional learning and memory.

#### Procedure

Participants completed two experimental sessions ∼10–12 hours apart. Day group participants were tested in the morning and again in the evening; night group participants were tested in the evening and returned the next morning after sleeping at home. Daytime wakefulness was selected as the comparison condition rather than sleep deprivation in order to isolate the contribution of natural sleep-dependent consolidation while minimizing additional physiological and stress-related confounds associated with sleep deprivation. In the first session (∼1 hr 15 min), participants completed sleepiness (SSS) and vigilance (PVT) assessments (∼10 min), followed by two discrimination blocks (∼25 min each) separated by a learning task (∼15 min). In the second session (∼1 hr 15 min), they completed another SSS and PVT (∼10 min) and a final discrimination task (∼25 min) before receiving monetary compensation. The experiment sessions started at 8:00–9:30 a.m. or 8:00–9:30 p.m., depending on group assignment. Sleep in the night group and wake activity in the day group were monitored via actigraphy between the sessions. (See Figure 1D, Supplementary Figure 2A).

### fMRI Experiment

#### Participants

Ninety healthy, right-handed participants were recruited for the fMRI experiment. They were assigned to a **day group** (n=45) or a **night group** (n=45). Written informed consent was obtained, and all procedures were approved by the IRB of the Weizmann Institute of Science and the Medical IRB of the Tel-Aviv Sourasky Medical Center. Ten participants were excluded due to technical issues with the scanner or task scripts (n=5), falling asleep during scanning (n=1), voluntary withdrawal (n=3), or excessive motion inside the scanner (n=1), and their data were neither preprocessed nor analyzed. The final analysis included 40 participants in the night group (21 females, mean age=26.97, SD=5.66) and 40 in the day group (25 females, mean age=26.1, SD=3.71). The fMRI experiment was designed following completion of the behavioral experiment and aimed to identify neural correlates of the behavioral sleep-related generalization effect rather than to test a specific ROI effect. Accordingly, the behavioral findings served as the empirical basis motivating the target sample size.

#### Procedure

Each participant completed two sessions separated by an approximately 10-hour consolidation interval. The day group was tested in the morning (the session started between 7:00–9:00 a.m.) and returned that evening; the night group was tested in the evening (the session started between 7:00–9:00 p.m.) and returned the following morning. The first session (∼2.5 hrs) included sleepiness and vigilance assessments (∼10 min), the first discrimination task block (∼25 min), followed by the learning task, which was conducted inside the MRI scanner (∼1 hour + preparation). After scanning, participants completed a second discrimination block (∼25 min) and left the lab. The second session matched the behavioral format, including SSS, PVT, and the final discrimination block. All discrimination blocks were performed outside the scanner. Sleep behavior in the night group and wake behavior in the day group were monitored using an actigraph watch between the sessions. (See Figure 1D and Supplementary Figure 2B).

### Sleep EEG Experiment

#### Participants

Forty-five healthy participants were recruited for the sleep EEG experiment. Written informed consent was obtained, and procedures were approved by the Medical IRB of the Tel Aviv Sourasky Medical Center. Three participants were excluded due to noisy or corrupted EEG data, resulting in a final sample of 42 participants (26 females, mean age=25.71, SD=3.17). Sample size was determined based on the logistical demands of overnight high-density EEG recordings and was comparable to previous sleep physiology studies conducted in the laboratory.

#### Procedure

All participants were assigned to a **night group** and slept in the lab under full polysomnographic and high-density EEG monitoring. They arrived ∼3 hours before their regular bedtime. Setup included EEG net application (∼1 hr), followed by SSS and PVT assessments (∼10 min), and two discrimination task blocks (∼25 min each) separated by a learning task (∼15 min), totaling ∼2.5 hours. After overnight sleep and a short recovery period, participants completed a second sleepiness and vigilance assessment (∼10 min), and a final discrimination block (∼25 min). The EEG net was removed, and participants were compensated. (See Figure 1D and Supplementary figure 2C).

## Method details

### Experimental tasks

#### Face stimuli used in the tasks

Twenty-six participants (14 women) rated faces with neutral expression sourced from an online database (the siblings DB)^70^, arranging them along a continuum from most malicious to most kind. Three males with similar average ratings as well as similar skin tones, hair, and eye colors were selected as the original faces (Figure 1A). These faces were positioned as vertices of a triangle space comprising 66 faces that were morphed composites of the three original (vertex) faces (FantaMorph software). The proximity of each face to each vertex corresponded to the proportion of the original face in the morph. Thus, the face positioned in the center of the triangle was an average representation derived from all three original faces (Figure 1A). The assignment of original faces to negative, positive, or neutral valence vertices was counterbalanced across participants. Twenty-nine (18 women) participants completed an online discrimination experiment to ensure no bias in perceiving morphed faces as similar to the original faces (see also ’Discrimination Task’ below).

#### Learning task

We used a probabilistic learning task (two-armed bandit^71^) to associate valence with the original face stimuli. We chose a probabilistic instrumental learning task rather than a Pavlovian design in order to examine how emotional generalization emerges through active reinforcement-based learning and decision-making, which more closely resembles naturalistic learning conditions. After a short training to ensure understanding, participants were instructed to choose one face in each trial and maximize their points according to feedback. During each trial, participants were presented with two out of three original faces simultaneously on the screen, with the left and right positions randomized for each trial. All possible pairs of faces were presented throughout the task. Each original face was probabilistically assigned (p=0.9) to either positive, negative, or neutral feedback (gain +10, loss -10, or zero points), and participants learned these associations as the task progressed. After their selection, the chosen face was highlighted by a yellow frame on the screen. Feedback regarding the participant’s response (gain +10, loss -10, or zero points) was then provided based on the selected face (see Figure 1B and Supplementary Figure 1A-C). Because face–feedback associations were highly predictable (p=0.9), participants quickly learned the value of each face and consistently chose the more rewarding option. This might lead to disproportionate exposure to positive feedback and limited exposure to negative feedback. To overcome this potential bias, we included ’helpless’ trials in which both presented stimuli were identical faces associated with negative or neutral feedback, ensuring balanced exposure across feedback types. Trial composition included a total of 132 trials comprising 30 positive-negative, 30 positive-neutral, 30 negative-neutral, 18 neutral-neutral, and 24 negative-negative face pairs trial.

In the fMRI experiment, fixation crosses appeared both between face selection and feedback, and during intervals between trials. The duration of the fixation cross was jittered uniformly between 1 and 7 seconds, with a mean of 4 seconds (Supplementary Figure 1B). The task lasted approximately one hour for the fMRI experiment, and included a brief pause of up to 2 minutes midway through, and 15 minutes for both the behavioral and sleep EEG experiments. In the sleep EEG experiment, participants first viewed two faces; after 1500 ms, arrows appeared beneath the faces, cueing participants to make a choice (Supplementary Figure 1C).

#### Discrimination task

The discrimination task operationalized generalization as the perceived similarity between images within the face space and each one of the original vertex faces. This task was performed in three blocks at different time points throughout the experiment: pre-learning, post-learning, and post-sleep/wake consolidation (Figure 1D, Supplementary Figure 2). At the beginning of the first discrimination block, participants received instructions and practiced the task in a short training. In the behavioral and fMRI experiments, each trial in the discrimination task begins with the presentation of a single morphed face image from within the face space. Then, participants are asked to choose one of the three original faces that most closely resembles the presented morphed face (Figure 1C, Supplementary Figure 1D, Supplementary Figure 2). Participants indicated their choice by pressing one of three arrows. As a result, the same screen was shown with a yellow rectangle surrounding the chosen face. The same trial was repeated if participants didn’t indicate a choice within three seconds. The correct choice earned participants points, while the incorrect choice resulted in a loss of points. Feedback was not provided after each trial; instead, the total score was presented at the end of each block. The accumulated points from all blocks of the discrimination task and the points earned in the learning task were translated into a monetary bonus at the end of the experiment.

In the EEG version of the experiment, participants initially had the opportunity to observe three original faces and memorize their appearance before the trials began. Subsequently, in each trial, the three original faces are presented briefly, followed by the appearance of the morphed face (random order). After 1000 ms (while the morph is still visible), arrows appear on the screen, allowing participants to make their selections (refer to Supplementary Figure 1E). In all experiments, each morph was presented six times (a total of 396 trials). The task lasted approximately 25 minutes in all experiments. (See Figure 1C, Supplementary Figure 1D, E and Supplementary Figure 2).

The modified temporal structure in both tasks in the EEG experiment was implemented to enable future analyses of task-related wake EEG dynamics. Although such analyses are beyond the scope of the present report and are not presented here, the timing was optimized to allow examination of stimulus-locked and decision-related neural activity.

#### Psychomotor Vigilance Task (PVT)

Participants were instructed to press a mouse button when numbers (representing a running RT timer) appeared on the screen. The inter-stimulus interval (ISI) varied randomly between 2–10 seconds, and the task lasted five minutes^72^. Sustained attention performance was assessed using mean and SD of RT, 1/RT, as well as lapse counts (RT > 500 ms) and errors (button presses without a stimulus). These metrics are highly sensitive to sleepiness, making the PVT a standard tool for vigilance-related assessments ^27,73,74^.

### Actigraphy

In the fMRI and behavioral experiments, we provided the participants with an actigraph watch (Ambulatory Monitoring, Inc. motionlogger Micro Watch). This watch measured acceleration, light, and temperature. These measures were used to validate sufficient sleep in the night groups and the absence of daytime naps in the day groups.

### fMRI

#### fMRI acquisition

Functional imaging data were collected using a 3T Siemens MAGNETOM Tim-Trio scanner with a 32-channel head coil. T2*-weighted images were acquired with a gradient-echo EPI sequence (TR=1450 ms, TE=30 ms, flip angle=90°), with 60 descending slices (3 mm thick, 10% gap) at a 30° coronal angle from the ACPC plane. Voxel size was 2.3 mm³ (FOV=220), and phase encoding was anterior-to-posterior. Functional scanning was conducted in two consecutive runs within the learning task, separated by a brief pause of up to 2 minutes. High-resolution anatomical T1-weighted images were also acquired (TR=2300 ms, TE=2.29 ms, flip angle=8°, voxel size=0.9 mm³, FOV=240). Functional scans covered nearly the entire brain, excluding a small dorsal parietal region. The anatomical scan covered the whole brain.

#### fMRI preprocessing

Preprocessing was performed using *fMRIPrep* 21.0.0 ^75,76^, based on Nipype 1.6.1 framework ^77,78^. Susceptibility distortion correction was performed using FSL’s TOPUP based on spin-echo EPI fieldmaps acquired with opposite phase-encoding directions (AP/PA). The estimated B0 field was subsequently applied to correct geometric distortions in the functional BOLD images^79^. T1-weighted anatomical images were corrected for intensity nonuniformity (N4BiasFieldCorrection^80^ implemented in ANTs version 2.3.3.^81^), skull-stripped (Nipype implementation of ANTs-based brain extraction using the OASIS30ANTs template), and segmented into gray matter (GM), white matter (WM), and cerebrospinal fluid (CSF) using FSL’s FAST^82^. Nonlinear spatial normalization to a standard space (MNI152NLin2009cAsym^83^) was completed using ANTs (antsRegistration). For each BOLD run, a reference volume was generated for motion correction (FSL mcflirt ^84^). Slice timing correction was applied (AFNI’s 3dTshift ^85^). Co-registration to the T1-weighted image was done using FreeSurfer’s *bbregister* with boundary-based registration ^86^. Functional images were resampled to MNI space using antsApplyTransforms (with Lanczos interpolation to minimize smoothing). Functional images were resampled to standard space using *antsApplyTransforms* with Lanczos interpolation, and surface-based resampling was done with FreeSurfer’s *mri_vol2surf*. Confound regressors included framewise displacement^84,87^, DVARS, and global signals (CSF, WM, whole brain). Temporal and anatomical CompCor were computed^88^ temporal (tCompCor) components were derived from the top 2% most variable voxels, and anatomical (aCompCor) components were extracted from masks of CSF, WM, and combined compartments, with GM faction minimized by excluding voxels with >5% GM partial volume ^88^. For each CompCor method, components explaining 50% of nuisance variance were retained. All regressors, including motion estimates and their derivatives/quadratics, were included in the confound file. Volumes exceeding FD > 0.9 mm or DVARS > 1.5 SD were flagged as motion outliers.

#### Whole-brain GLM analysis

Voxel-wise statistical analyses were conducted across the whole brain. At the first level, individual events were modeled using a canonical hemodynamic response function (HRF), and framewise displacement estimates from preprocessing were included as nuisance regressors to control for motion. Each participant’s two functional runs were modeled separately using a first-level general linear model (GLM), yielding subject-specific statistical maps for each modeled event type. The event-related design included six regressors, corresponding to the appearance of the face’s stimulus and the feedback, each with positive, negative, or neutral valence. For the face stimuli, valence was determined by the expected feedback associated with the participant’s choice. Regressors were time-locked to event onset and modeled over the full event duration: three seconds for the stimulus and two seconds for the feedback. Subject-level statistical maps were subsequently entered into a second-level group analysis. All participants were included in the group-level model regardless of sleep condition (day/night), enabling unbiased voxel selection for downstream analysis. The group-level model was implemented as an intercept-only (one-sample) analysis across all participants. The resulting voxelwise statistics were converted to equivalent Z-scores, and statistical thresholding was performed on these Z-score maps. Group-level statistical maps were corrected for multiple comparisons using a family-wise error rate (FWER) threshold of p < 0.001, with a minimum cluster size of 10 voxels. These group-level maps were used as functional localizers for ROI selection and peak-voxel identification and were independent of the subsequent behavioral generalization analyses.

#### ROI selection

A predefined list of 28 regions of interest (ROIs) was compiled based on the Harvard-Oxford Atlas, guided by prior research implicating these regions in reward/punishment processing, decision-making, emotion regulation, and face perception. ROI selection for downstream analyses was based specifically on activation during the feedback phase, as the later analyses focused on neural responses associated with emotional reinforcement processing. ROIs were included in the final analysis only if they belonged to the predefined ROI list and showed significant task related activation in the group level phase. Fourteen regions met these criteria. See Table 1.

**Table 1:** List of 28 a priori regions of interest (ROIs) based on the Harvard-Oxford Atlas. Fourteen ROIs that passed statistical thresholding (FWE-corrected p < 0.001, cluster size ≥ 10 voxels) were included in the final analysis (left column).

| **ROI name** | **Included?** | **ROI name** | **Included?** |
|---|---|---|---|
| Caudate | V | Cuneus | X |
| Putamen | V | Thalamus | X |
| Pallidum | V | Posterior Superior temporal gyrus | X |
| Hippocampus | V | Occipital Fusiform Gyrus | X |
| Amygdala | V | Fusiform Face Area | X |
| Accumbens | V | Occipital Face Area | X |
| Precuneus | V | Posterior Temporal Fusiform Cortex | X |
| Orbitofrontal Cortex | V | Anterior Temporal Fusiform Cortex | X |
| Medial Frontal Cortex | V | Temporo-occipital fusiform cortex | X |
| Subcallosal Cortex | V | Superior Temporal Sulcus | X |
| Anterior Cingulate Cortex | V | Anterior Middle Temporal Gyrus | X |
| Posterior Cingulate Gyrus | V | Posterior Middle temporal Gyrus | X |
| Anterior Superior Temporal Gyrus | V | Postcentral Gyrus and the Brainstem | X |
| Insula | V | Brainstem | X |

For each of the 14 ROIs, a 3 mm-radius sphere was defined around the peak activation voxel. Peak voxels were identified separately for each contrast from the group-level functional localizer maps within each anatomically defined ROI. This approach was chosen to obtain a localized and robust estimate of task-related activity while avoiding reliance on a single voxel. Peak-centered spherical ROIs provide a localized measure of activation while reducing the influence of functional heterogeneity that may exist within larger anatomical regions^89,90^. Given the functional voxel size (2.3-mm isotropic), a 3-mm radius corresponds to a small local neighborhood that preserves anatomical specificity while improving signal robustness. Activity was extracted from both the peak voxel and its surrounding sphere using each participant’s unthresholded statistical maps. If a sphere extended beyond anatomical ROI boundaries, only the portion within the ROI was retained.

To assess differences in neural responses, we computed two contrasts: Gain (positive > neutral feedback) and Loss (negative > neutral feedback). These contrasts were defined at the feedback stage rather than at stimulus onset, as our primary interest was in how neural responses to positive and negative reinforcement signals predict subsequent generalization and overgeneralization behavior. Contrast values were averaged across both scanning sessions for each participant and used in subsequent correlational analyses with behavioral generalization measures.

### Sleep EEG

#### Data Acquisition

Continuous high-density EEG and full polysomnography were recorded throughout the entire experimental session, including both cognitive tasks and sleep periods. A 256-channel hydrogel Geodesic Sensor Net (Electrical Geodesics, Inc., EGI) was used for data acquisition. Each electrode consisted of a silver chloride carbon fiber pellet, a lead wire, and a gold-plated pin, and was filled with conductive gel (Electro-Cap International). Signals were amplified using an AC-coupled, high-input-impedance amplifier (NetAmps 300, EGI) with an integrated antialiasing analog filter. Recordings were referenced to Cz and digitized at a sampling rate of 1000 Hz. Prior to recording, electrode impedance was verified to be below 50 kΩ for all sensors.

#### Sleep Scoring

Sleep stages were manually scored offline in accordance with the American Academy of Sleep Medicine (AASM) guidelines. Each recording was scored by two trained raters to ensure reliability. Standard sleep parameters, including sleep latency, total sleep time, and time spent in each sleep stage, were calculated for subsequent analysis. To examine whether REM fragmentation was associated with overnight generalization, we quantified REM continuity and transient high-frequency arousals during REM sleep using an approach adapted from Wassing et al. (2019)^91^. Briefly, transient cortical arousals were identified as events with spectral activity above 16 Hz lasting 3–15 s and thresholded relative to each participant’s N3 baseline activity. For each participant, we calculated REM arousal counts and REM arousal density (number of arousals divided by total REM duration). These measures were subsequently tested for associations with overnight generalization outcomes.

#### Preprocessing

Analyses were conducted in Python using the sleepEEGpy package^36^, an open-source "wrapper" pipeline that integrates MNE-Python^92^, PyPREP^93^, YASA^37^, SpecParam^94^, and other tools. Each EEG recording was down-sampled from 1000 Hz to 250 Hz and band-pass filtered between 0.75–40 Hz using a zero-phase FIR filter with a Hamming window. Manual inspection of raw signals was performed using the "butterfly" view, complemented by topographical maps to identify and exclude abnormal channels. Subsequently, Independent Component Analysis (ICA) was applied using the extended infomax algorithm^95^ with 40 Principal Component Analysis (PCA) components. Components associated with ECG artifacts were identified through visual inspection and removed.

#### Power Spectral Analysis

Power Spectral Density (PSD) was computed separately for each sleep stage using the Welch method as implemented in SleepEEGpy ^36^. Parameters included a Fast Fourier Transform (FFT) length of 2048 samples, a Hamming window, and a 50% segment overlap (1024 samples). Spectral power was quantified in the delta (0.75–4 Hz) and sigma (12.5–15 Hz) frequency bands during NREM sleep, and in the theta band (4–8 Hz) during REM sleep. For each participant and each frequency band, the channel with the maximum band-limited power across all electrodes was identified (see Supplementary Figure 9 for peak sigma band electrode distribution). The band power extracted from this peak electrode was used for subsequent correlation analyses.

#### Sleep Spindle Detection

Analyses were conducted in Python using sleepEEGpy^36^ employing the YASA algorithm ^37^. Spindles were detected during Non-rapid eye movement (NREM) sleep from each participant’s peak sigma channel. Fast spindles were defined within the 13–15 Hz frequency range, with durations constrained between 0.5–2.0 seconds and a minimum inter-spindle interval of 500ms. Detection criteria included: Correlation ≥ 0.65 (moving correlation between the raw and sigma-filtered signal), Relative power ≥ 0.2 (ratio of power in the spindle band to broadband power), root-mean-square (RMS) amplitude ≥ 1.5 standard deviations above the mean of the moving RMS of the sigma-filtered signal.

## Quantification and statistical analysis

### Generalization analysis

#### Generalization assessment

We analyzed each participant’s response patterns separately for each session. We calculated the proportion of responses classifying each morphed face as most similar to one of the original faces, organizing them into three triangular spaces based on valence: positive, negative, or neutral. As face–valence pairings were counterbalanced, triangles were labeled by valence rather than face identity (Figure 1E, top left). We interpolated each triangle to generate continuous response gradients (Figure 1E, top right). Perceptual boundaries were defined using a 50% threshold, such that a morphed face was considered to belong to a given original face if it was classified as resembling that face on at least 50% of its presentations. The area within each triangle meeting this criterion was defined as the generalization region. The size of these areas served as a measure of generalization (Figure 1E, bottom left; for visualization, all three are shown on one triangle). To assess changes over time, we computed two metrics: (1) immediate generalization: the area change from Block 1 to Block 2 (after learning), and (2) delayed generalization: the change from Block 2 to Block 3 (after sleep or wake; Figure 1E, bottom right). Note: While a 66% threshold was used in the figures for clearer visualization, all analyses were conducted using the 50% threshold.

#### Repeated face presentations

Although the discrimination task was repeated across three blocks, it could, in principle, introduce additional learning through repeated exposure to the faces, these trials were conducted without immediate feedback. Thus, repeated exposure would be expected to promote extinction or weakening of learned valence associations rather than reinforcement, potentially reducing the magnitude of generalization effects. Therefore, the observed generalization patterns are unlikely to be explained solely by repeated task exposure.

#### Generalization score

In the fMRI experiment, we computed a single score to capture the balance between generalization of positive and negative faces. This generalization score was calculated by subtracting negative face generalization from positive face generalization. Scores above zero indicate a bias toward generalizing the positive face, while scores below zero indicate greater generalization of the negative face. This measure reflects the idea that increased generalization of one emotional valence occurs at the expense of the other.

#### Renormalization index

To quantify changes in generalization over time, we computed a renormalization index for each participant by subtracting the immediate generalization score from the delayed generalization score. In the fMRI experiment, we used the generalization bias score (positive minus negative) for this calculation. In the EEG experiment, we focused on generalization of the positive face only.

#### Overgeneralization Calculation

Overgeneralization was defined as instances where participants incorrectly classified morphed faces as resembling one of the original faces, beyond the boundaries of the correct generalization zone. In the generalization task, such incorrect responses resulted in point loss and were considered maladaptive. Regions of the triangular face space that extended beyond the adaptive generalization boundaries were designated as overgeneralization areas. See Supplementary Figure 8A, B for illustration.

## Statistical analysis

Group-level effects in generalization were assessed using mixed-model ANOVAs. Significant interactions were followed by paired or independent-samples two-tailed t-tests, depending on the comparison, with correction for multiple comparisons using the Benjamini–Hochberg procedure. To examine whether brain ROI activity predicted generalization, we applied both linear regression and LASSO (least absolute shrinkage and selection operator) regression. All predictors were z-scored prior to LASSO to ensure comparability. The regularization parameter was optimized using 10-fold cross-validation and averaged across folds. Linear regression models were then constructed using the subset of predictors selected by LASSO.

Pearson correlations were used to assess associations between behavioral generalization scores and neural measures. Specifically, we examined correlations between amygdala activation (from ROI-based fMRI contrasts) and generalization indices, as well as between EEG-derived metrics (e.g., spindle count and sigma power) and changes in generalization across discrimination blocks. Statistical significance was set at *p* < 0.05.

To facilitate interpretation of both significant and non-significant findings, sensitivity analyses were conducted using the primary statistical model for each experiment. Assuming a two-sided significance threshold of α = .05 and 80% power, the smallest detectable effect for the behavioral Face × Group interaction corresponded to partial η² = .079 (Cohen’s f = .292). For fMRI ROI association analyses using the full sample (N = 80), the smallest detectable association was |r| = .306. For the sleep EEG experiment (N = 42), the smallest detectable correlation was |r| = .413. These analyses indicate that the present studies were primarily sensitive to effects in the moderate range, whereas smaller effects may not have been reliably detectable.

### Learning performance

Learning accuracy was quantified as the percentage of correct choices across all trials and during the final 10 trials. Learning trajectories were analyzed using trial-by-trial mixed-effects models with participant as a random effect. Group (sleep/wake) and face-pair effects were evaluated using mixed-effects models and FDR-corrected pairwise comparisons.

### Discrimination Accuracy

Discrimination accuracy was calculated separately for each participant and session as the percentage of correct responses among trials receiving a correct or incorrect response. Equal-blend/no-answer trials were counted separately and excluded from the accuracy denominator. Session and group effects were assessed using linear mixed-effects models with participant-level random intercepts and by-participant random slopes for Session, allowing the effect of Session to vary across participants. Session was treated as a categorical within-participant factor. For experiments including both day and night groups, the model included fixed effects of Session, Group, and the Session × Group interaction (`accuracy ∼ Session * Group + (1 + Session | participant)`). For the EEG experiment, which included a single group, the model included Session as the fixed effect (`accuracy ∼ Session + (1 + Session | participant)`). Day–night differences within each session were additionally assessed using Welch’s independent-samples t-tests, with Benjamini–Hochberg correction across sessions.

To assess whether sleepiness and vigilance influenced the fMRI findings, we fitted a Gaussian generalized estimating equation with an exchangeable within-subject correlation structure. Face condition, group, and their interaction were included as predictors, with centered session-2 SSS and PVT mean reaction time as covariates. Data were matched by subject ID, and participants missing either covariate were excluded.

### Figure significance markers

In the Figures, levels of significance were denoted by **/*/∼, respectively corresponding to p-values <0.01/<0.05</0.10. The significance marker, together with the name of the test used, are located at appropriate places in the Figure legends.

## Learning task analysis

To confirm that participants learned the associations between faces and outcomes, we calculated the percentage of correct choices for each participant across all trials. In all experiments, all participants performed above chance level, with a minimum accuracy of 60%. The average accuracy across all three experiments was 93.1% (SD=5.9%). The learning curve shown in Figure 1B reflects data from all participants across the three experiments (N=182), smoothed using a moving average over a window of five trials.

## Key resources table

| REAGENT or RESOURCE | SOURCE | IDENTIFIER |
|---|---|---|
| Deposited data |  |  |
| Raw and behavioral data from all three experiments | Mendeley Data | DOI: 10.17632/s9vxcsk9yx.1 |
| Software and algorithms |  |  |
| FantaMorph | Abrosoft |  |
| fMRIPrep | Esteban et al.; fMRIPrep developers | Version 21.0.0 |
| Nipype | Gorgolewski et al. | Version 1.6.1 |
| FSL | FMRIB, University of Oxford | Version 6.0.5.1 |
| ANTs | Advanced Normalization Tools | Version 2.3.3 |
| FreeSurfer | Martinos Center for Biomedical Imaging | Version 6.0.1 |
| AFNI | NIH |  |
| Python | Python Software Foundation | Versions: 3.10.16, 3.13.2, 3.11.15, 3.10.6, 3.9.21, 3.9.7 |
| sleepEEGpy | Falach et al., 2025 | DOI: 10.1016/j.compbiomed.2025.110232; Zenodo: 10.5281/zenodo.10362189 |
| MNE-Python | Gramfort et al., 2013 | DOI: 10.3389/fnins.2013.00267; Zenodo: 10.5281/zenodo.592483 |
| PyPREP | Bigdely-Shamlo et al., 2015 / PyPREP developers | DOI: 10.3389/fninf.2015.00016; Zenodo: 10.5281/zenodo.6363575 |
| YASA | Vallat and Walker, 2021 | DOI: 10.7554/eLife.70092 |
| SpecParam | Donoghue et al., 2020 | DOI: 10.1038/s41593-020-00744-x |
| MATLAB | MathWorks | Version 2022a; RRID: SCR_001622 |

## Acknowledgments

We thank Eyal Hizmi and Maya Klinovsky for their assistance with data collection; Nofar Shemen, Eden Zohar, and Adit Friedberg for their assistance with data collection and preliminary analyses; Rotem Falach for her assistance with EEG analysis. We also thank Shir Frank, Gal Zatelman, Tomer Cohen, Rakefet Chishinski, Reut Rittal, and Hadar Nakar for their assistance with EEG data collection and sleep scoring. We are grateful to Liliya Iskhakova, Tamir Scherf, Revital Shani-Hershkovich, Noa Bar-Ilan, Sasha Devore and others for their support and assistance.

This work was supported by grants from the Israel Science Foundation (ISF #1467/24) and the European Research Council (ERC-2023-ADG #101142391) granted to Rony Paz, and by grants from the Israel Science Foundation (ISF #1557/22) and the European Research Council (ERC-2019-CoG #864353) to Yuval Nir. Additionally, the Nir lab is supported by the Sieratzki Institute for Advances in Neurosciences.

## Author Contributions

**Conceptualization:** Ella Bar, Rony Paz, Yuval Nir;

**Methodology:** Ella Bar, Rony Paz, Yuval Nir;

**Investigation:** Ella Bar, Max Bringman, Reut Weissgross, Kristoffer Aberg;

**Formal Analysis:** Ella Bar, Max Bringman;

**Software:** Ella Bar, Max Bringman, Gennadiy Belonosov;

**Data Curation:** Ella Bar;

**Visualization:** Ella Bar;

**Writing – Original Draft:** Ella Bar;

**Writing – Review & Editing:** Ella Bar, Rony Paz, Yuval Nir;

**Supervision:** Rony Paz, Yuval Nir;

**Funding Acquisition:** Rony Paz, Yuval Nir.

