## Supplementary Material for "Sleep Renormalizes Negative Emotional Generalization"

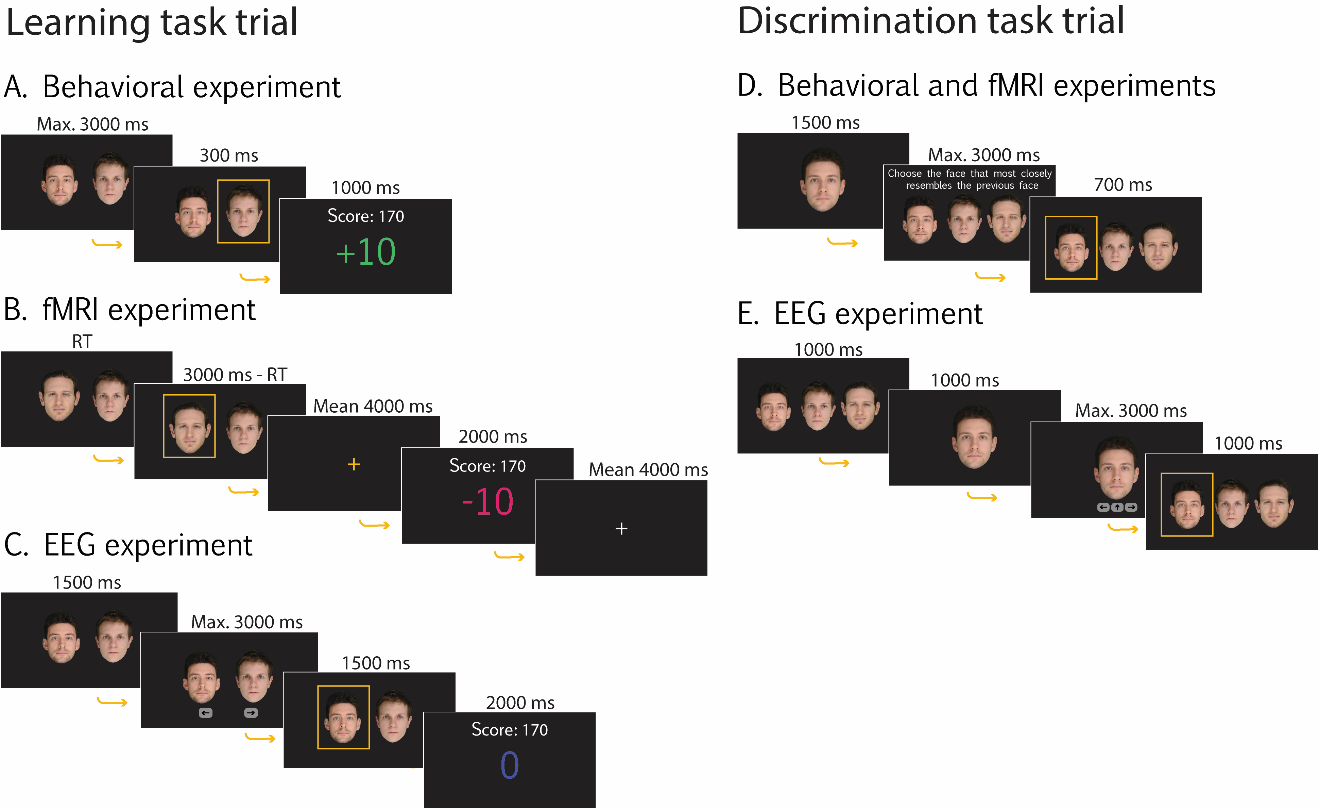

**Supplementary Figure 1. Trial timelines for the learning and discrimination tasks across experiments. Related to Figure 1 and STAR Methods.**

**A–C.** Timeline of learning task trials across experiments. Feedback was either +10, -10, or 0 points based on participants’ responses. **A.** Behavioral experiment trial structure. **B.** fMRI experiment trial structure. A fixation cross appeared before feedback and between trials, with durations jittered uniformly between 1–7 seconds (mean=4 seconds). **C.** EEG experiment trial structure. Choices could only be made after arrows appeared on the screen. **D, E.** Timeline of discrimination task trials**. D.** Behavioral and fMRI experiments. **E.** EEG experiment. Choices were permitted only after the arrow's onset.

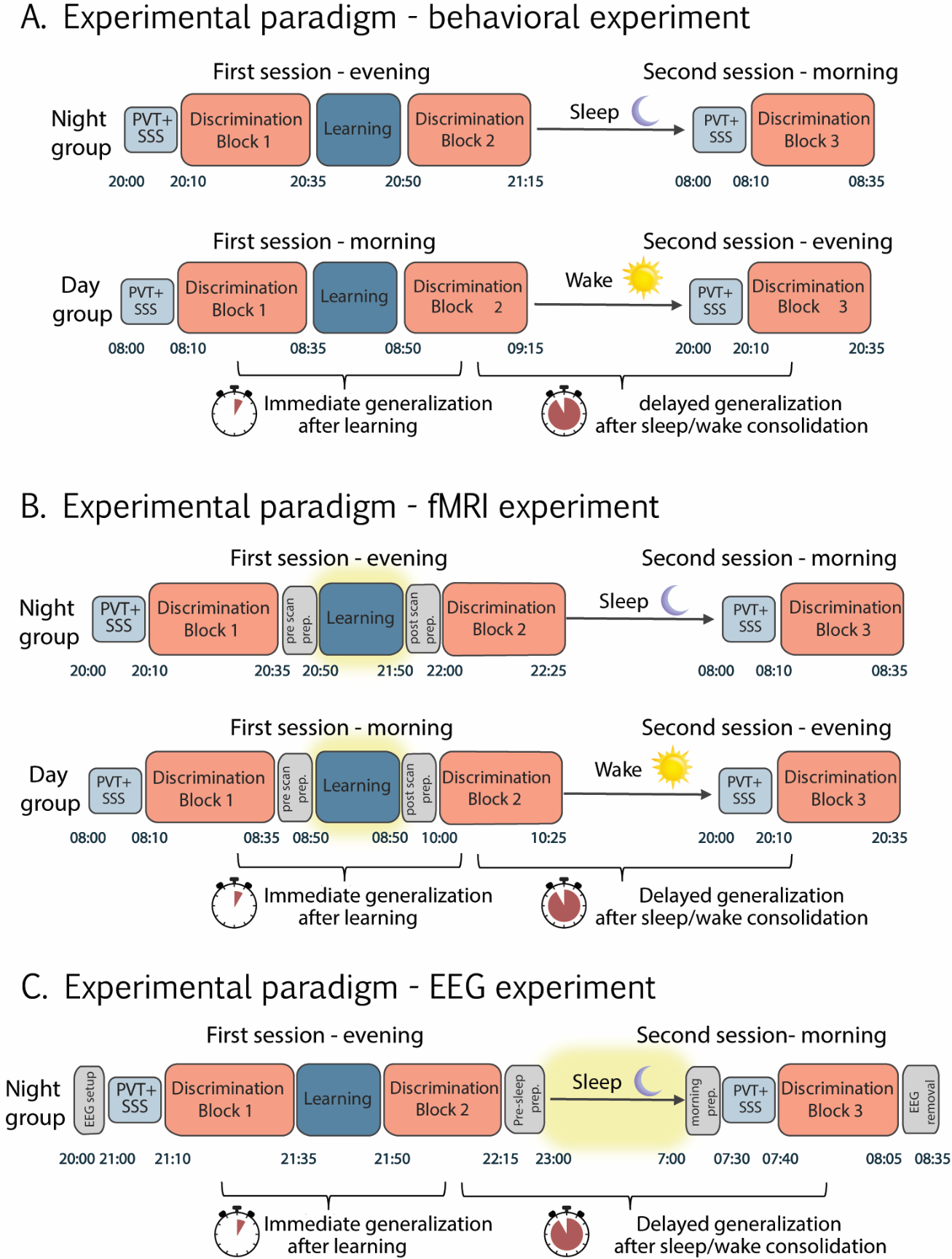

**Supplementary Figure 2. Experimental paradigms across behavioral, fMRI, and EEG experiments. Related to Figure 1 and STAR Methods.**

**A. Behavioral experiment. Participants were randomly assigned to either a night group or a day group and completed two experimental sessions. In the night group, the first session began in the evening (start time between 19:00 and 21:00), whereas in the day group it began in the morning (start time between 07:00 and 09:00). Each session began with the Psychomotor Vigilance Task (PVT) and the Subjective Sleepiness Scale (SSS) to control for fatigue and time-of-day effects. During the first session, following the PVT and SSS, participants completed two discrimination blocks with a reinforcement learning phase in between. The two sessions were separated by a 10–12 h consolidation interval consisting of either overnight sleep (night group) or daytime wakefulness (day group), during which participants wore an actigraphy watch to monitor sleep and activity. The second session included a third and final discrimination block. Immediate generalization was defined as the change from discrimination block 1 to block 2, and delayed generalization as the change from block 2 to block 3. Time stamps indicate the approximate start time of each session for a representative participant. B. fMRI experiment. The paradigm was identical to the behavioral experiment, except that the learning phase was performed inside the MRI scanner and was longer due to extended inter-stimulus (ISI) and inter-trial (ITI) intervals, resulting in a longer session that included pre- and post-scan preparation as well as scanning time (timing is indicated in the figure for a representative participant). The scan period is indicated by a yellow frame. Discrimination blocks were performed outside the scanner. As in A, the two sessions were separated by a 10–12 h consolidation interval consisting of either overnight sleep (night group) or daytime wakefulness (day group), during which participants wore an actigraphy watch to monitor sleep and activity. Group structure (night vs. day) and definitions of immediate and delayed generalization were the same as in A.** **C. EEG experiment. The paradigm was similar to the behavioral experiment, with the exception that only a night group was included. High-density EEG was recorded continuously throughout the experiment; however, only sleep data during the consolidation interval between the two sessions were analyzed, and this period is indicated by a yellow highlighted frame in the figure. The recording session included EEG setup prior to sleep and morning procedures following awakening (timing is indicated in the figure for a representative participant). Task structure was identical to (A). Definitions of immediate and delayed generalization were the same as in (A).**

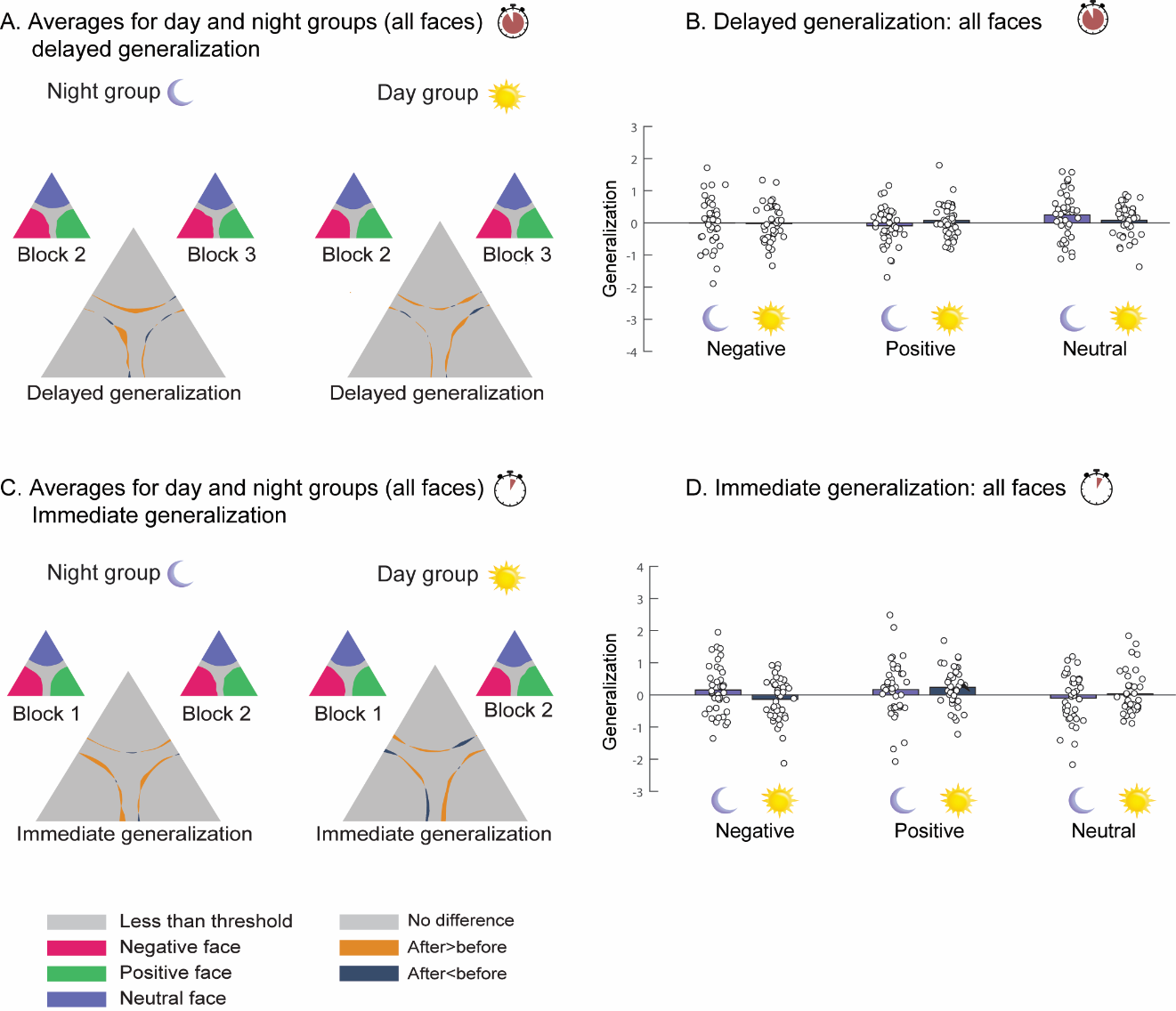

**Supplementary Figure 3. No significant group differences between the day and night groups in the fMRI experiment. Related to Figure 2.**

**A.** Group-level delayed generalization: Small triangles represent average morph areas classified as each of the original faces during discrimination blocks 2 and 3, overlaid onto a single triangle. Large triangles depict the change in generalization between blocks. Left: night group; right: day group. No significant group differences were observed. **B.** Summary of delayed generalization: Bar graph shows the average change in generalization for positive, negative, and neutral faces. No significant differences between groups.
**C.** Group-level immediate generalization: Same format as (A), comparing discrimination blocks 1 and 2. No significant changes in generalization for either group. **D.** Summary of immediate generalization: Bar graph shows average generalization change for all faces. No significant group differences were observed.

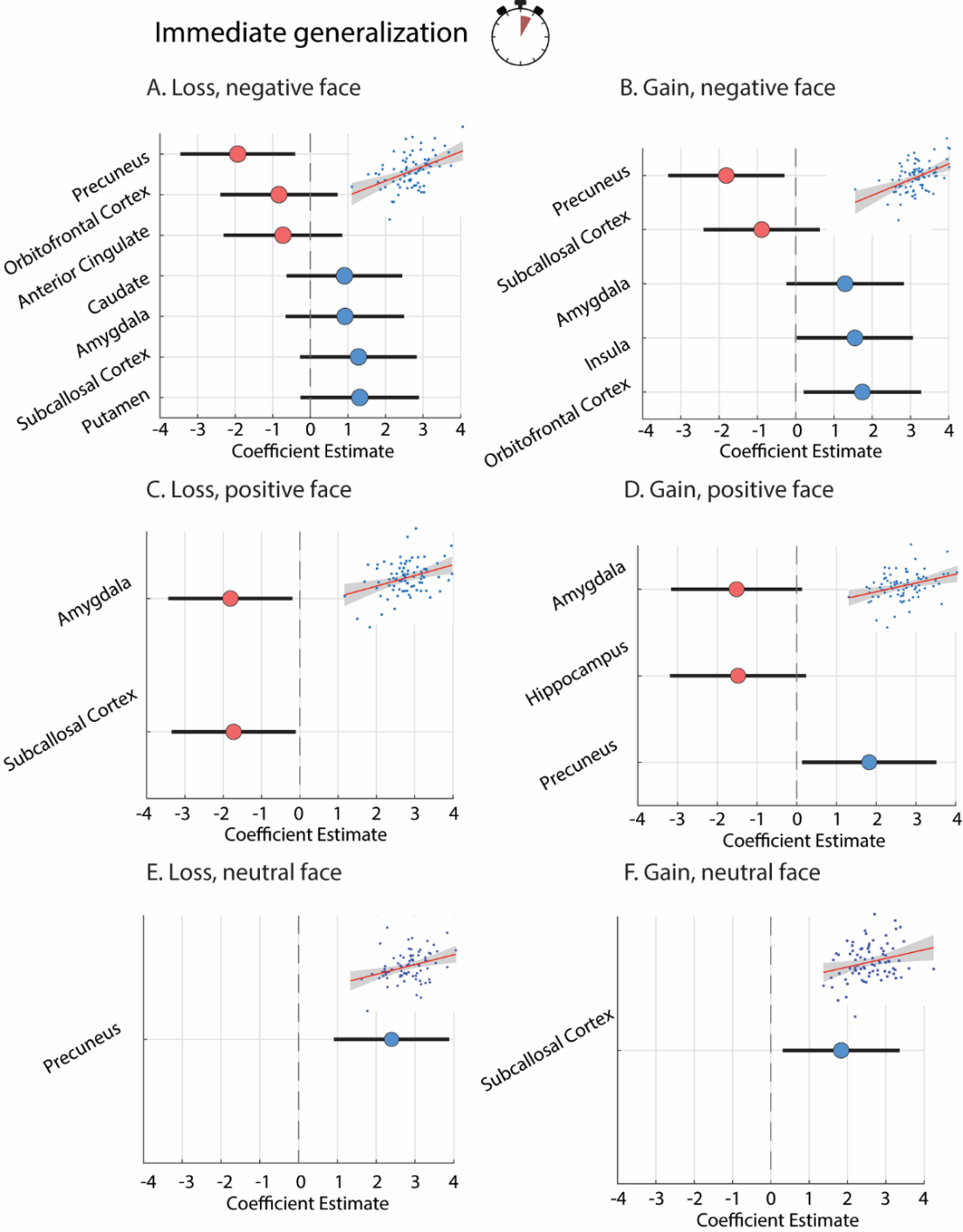

**Supplementary Figure 4. Neural activity predicts immediate generalization.Related to Figure 3.**

**A–F.** Lasso regression models predicting immediate generalization of negative, positive, and neutral faces from ROI activity during learning, separately for loss and gain outcomes. Insets show model fits (x-axis: predicted; y-axis: observed; red: fit; gray: 95% CI). Positive coefficients indicate that higher activity was associated with greater generalization, whereas negative coefficients indicate reduced generalization. **A–B.** Negative face generalization. **A.** Loss: Higher activity in the Caudate, Amygdala, Subcallosal Cortex, and Putamen was associated with greater negative-face generalization, whereas activity in the OFC, ACC, and Precuneus was associated with reduced generalization (R²=0.21, p=0.012). **B.** Gain: Activity in the Amygdala, Insula, and OFC was associated with greater negative-face generalization, while activity in the Subcallosal Cortex and Precuneus was associated with reduced generalization (R²=0.21, p=0.003). **C–D.** Positive face generalization. **C.** Loss: Activity in the Amygdala and Subcallosal Cortex was associated with reduced positive-face generalization (R²=0.12, p=0.007). **D.** Gain: Activity in the Amygdala and Hippocampus was associated with reduced positive-face generalization, whereas activity in the Precuneus was associated with increased generalization (R²=0.12, p=0.02). **E–F.** Neutral face generalization. **E.** Loss: Activity in the Precuneus was associated with increased neutral-face generalization (R²=0.12, p=0.002). **F.** Gain: Activity in the Subcallosal Cortex was associated with increased neutral-face generalization (R²=0.07, p=0.02).
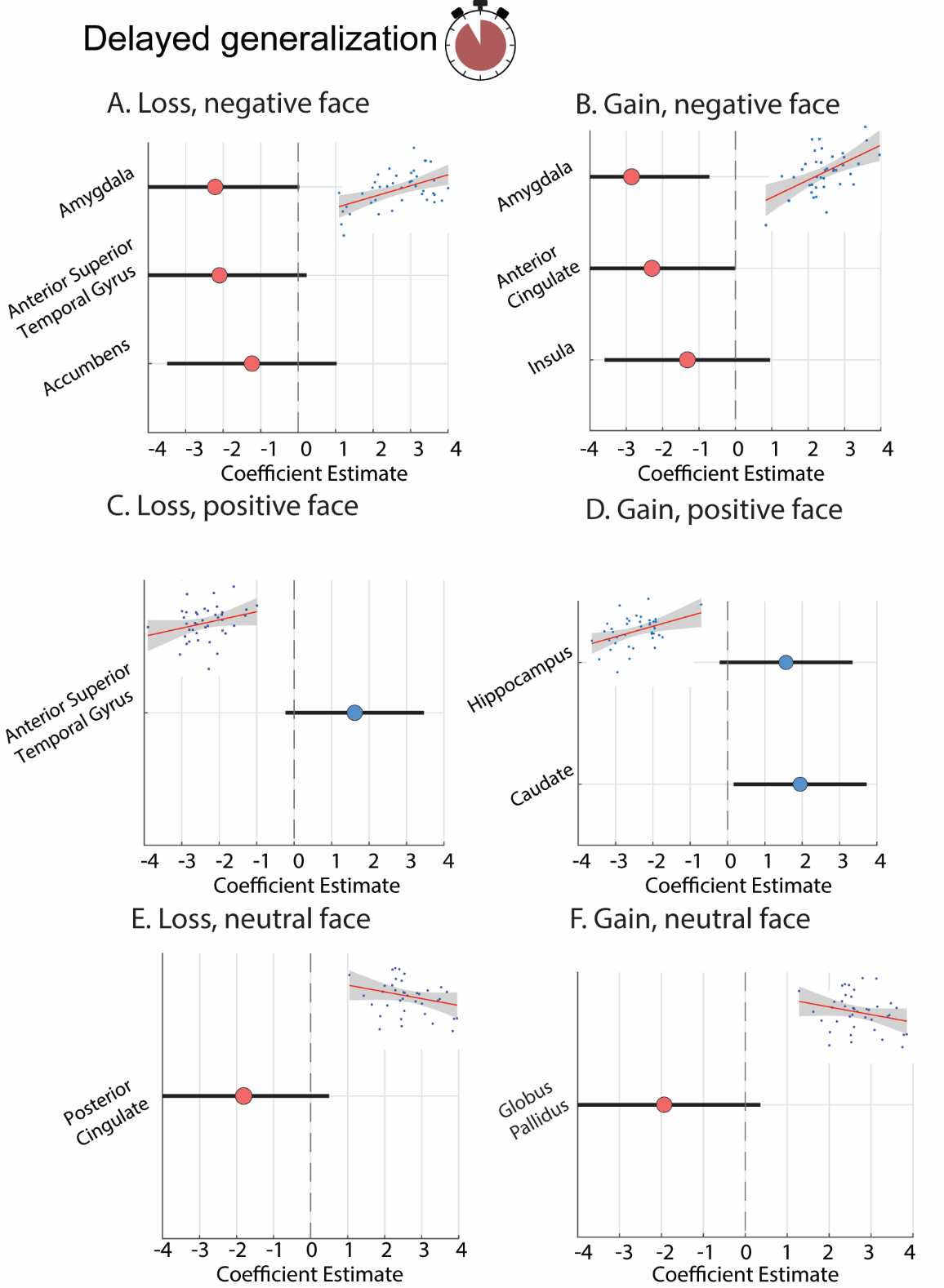

**Supplementary Figure 5. Neural activity predicts delayed generalization (Night group). Related to Figure 3.**

**A–F.** Lasso regression models predicting delayed generalization of negative, positive, and neutral faces from ROI activity during learning, separately for loss and gain outcomes. Insets show model fits (x-axis: predicted; y-axis: observed; red: fit; gray: 95% CI). Positive coefficients indicate that higher activity was associated with greater generalization, whereas negative coefficients indicate reduced generalization. **A–B.** Negative face generalization. **A.** Loss: Higher activity in the Amygdala, Anterior Superior Temporal Gyrus, and Accumbens was associated with reduced negative-face generalization (R²=0.26, p=0.013). **B.** Gain: Activity in the Amygdala, Anterior Cingulate Cortex, and Insula was associated with reduced negative-face generalization (R²=0.29, p=0.005). **C–D.** Positive face generalization. **C.** Loss: Activity in the Anterior Superior Temporal Gyrus was associated with increased positive-face generalization (R²=0.08, p=0.08). **D.** Gain: Activity in the Hippocampus and Caudate was associated with increased positive-face generalization (R²=0.17, p=0.03). **E–F.** Neutral face generalization. **E.** Loss: Activity in the Posterior Cingulate Cortex was associated with reduced neutral-face generalization (R²=0.06, p=0.12). **F.** Gain: Activity in the Globus Pallidus was associated with reduced neutral-face generalization (R²=0.07, p=0.09).

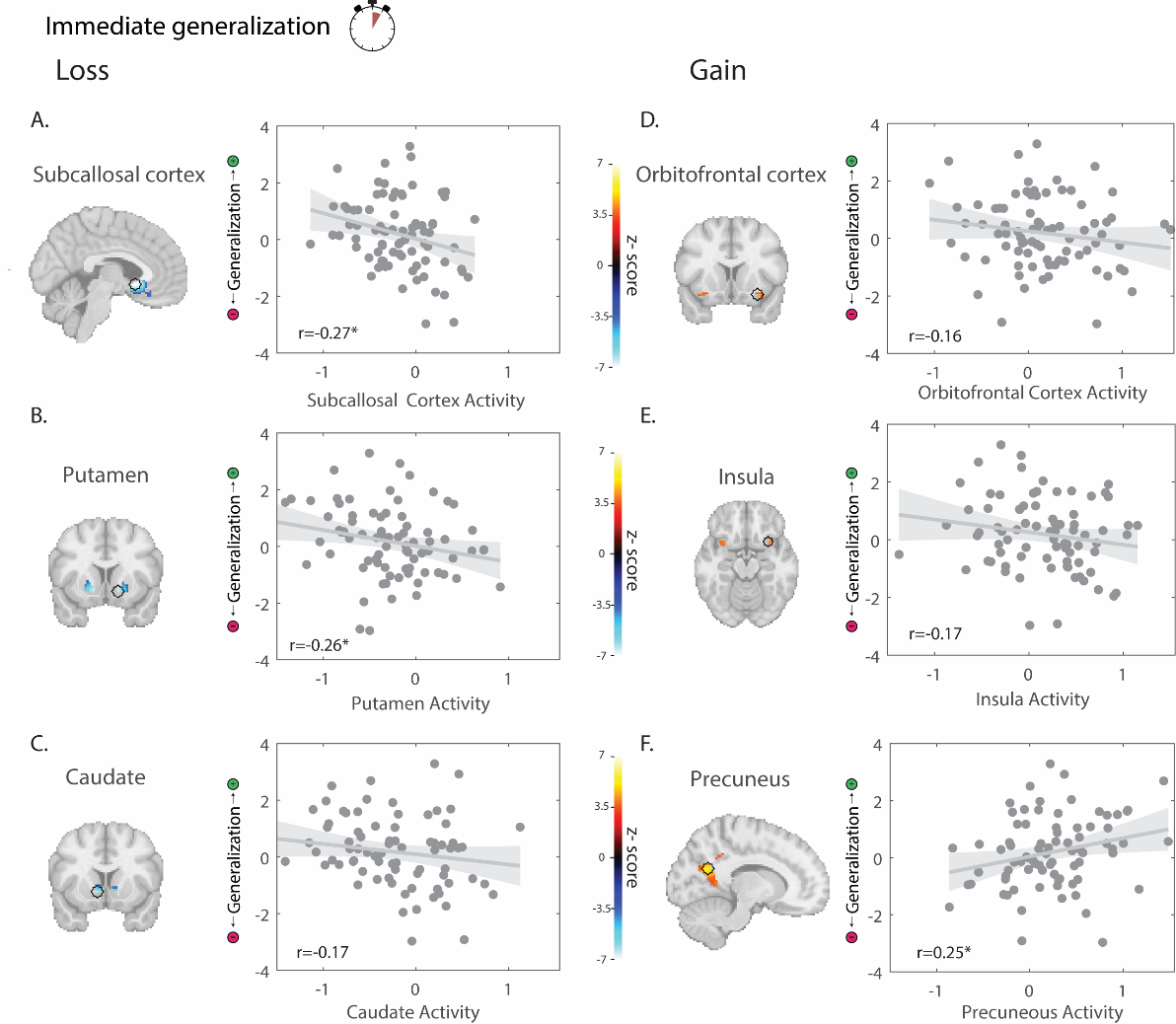

**Supplementary Figure 6. ROI activity correlates with immediate generalization. Related to Figures 3 and 4.**

**A–F.** Correlations between activity in ROIs selected via Lasso regression as predictors of immediate generalization, in addition to the Amygdala reported in the main text. Left panels show activation maps with a black circle representing a 3-mm-radius sphere centered on the peak voxel within each anatomical ROI (for visualization); analyses were based on mean activity within this sphere. Right panels show correlations between ROI activity during learning and immediate generalization. Negative correlations indicate that higher activity was associated with greater negative-face generalization (and reduced positive-face generalization), whereas positive correlations indicate greater positive-face generalization. **A.** Loss: Subcallosal Cortex activity was negatively correlated with immediate generalization (rₚ=–0.27, p=0.01). **B.** Loss: Putamen activity was negatively correlated with immediate generalization (rₚ=–0.26, p=0.02). **C.** Loss: Caudate activity showed a negative, non-significant correlation (rₚ=–0.17, p=0.12). **D.** Gain: OFC activity showed a negative, non-significant correlation (rₚ=–0.16, p=0.15). **E.** Gain: Insula activity showed a negative, non-significant correlation (rₚ=–0.17, p=0.12). **F.** Gain: Precuneus activity was positively correlated with immediate generalization (rₚ=0.25, p=0.02).

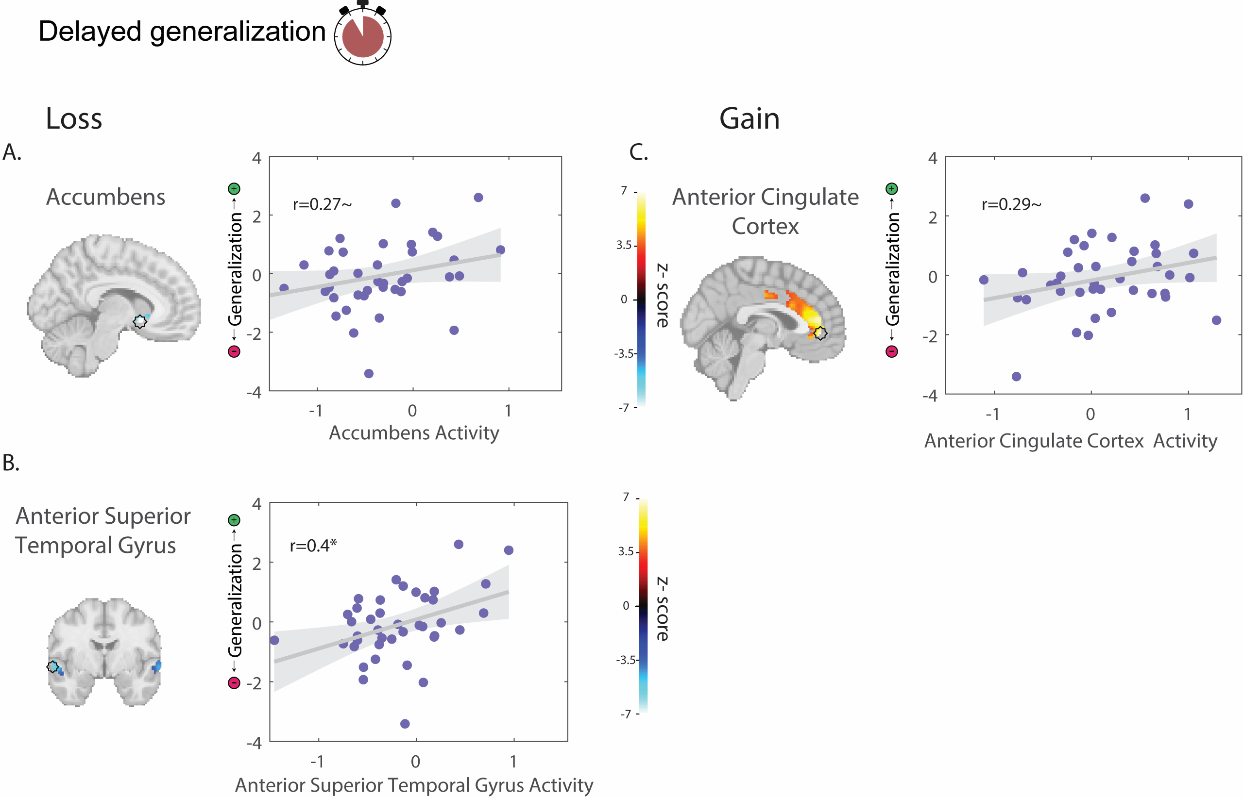

**Supplementary Figure 7. ROI activity correlates with delayed generalization (Night group). Related to Figures 3 and 4.**

**A–C.** Correlations between activity in ROIs selected via Lasso regression as predictors of delayed generalization after sleep, in addition to the Amygdala reported in the main text. Left panels show activation maps with a black circle representing a 3-mm-radius sphere centered on the peak voxel within each anatomical ROI (for visualization only); analyses were based on mean activity within this sphere. Right panels show correlations between ROI activity during learning and delayed generalization. Positive correlations indicate that higher activity was associated with greater positive-face generalization, whereas negative correlations indicate reduced positive-face generalization. **A.** Loss: Accumbens activity showed a positive, non-significant correlation with delayed generalization (rₚ=0.27, p=0.08). **B.** Loss: Anterior Superior Temporal Gyrus activity was positively correlated with delayed generalization (rₚ=0.40, p=0.01). **C.** Gain: Anterior Cingulate Cortex activity showed a positive, non-significant correlation with delayed generalization (rₚ=0.29, p=0.07).

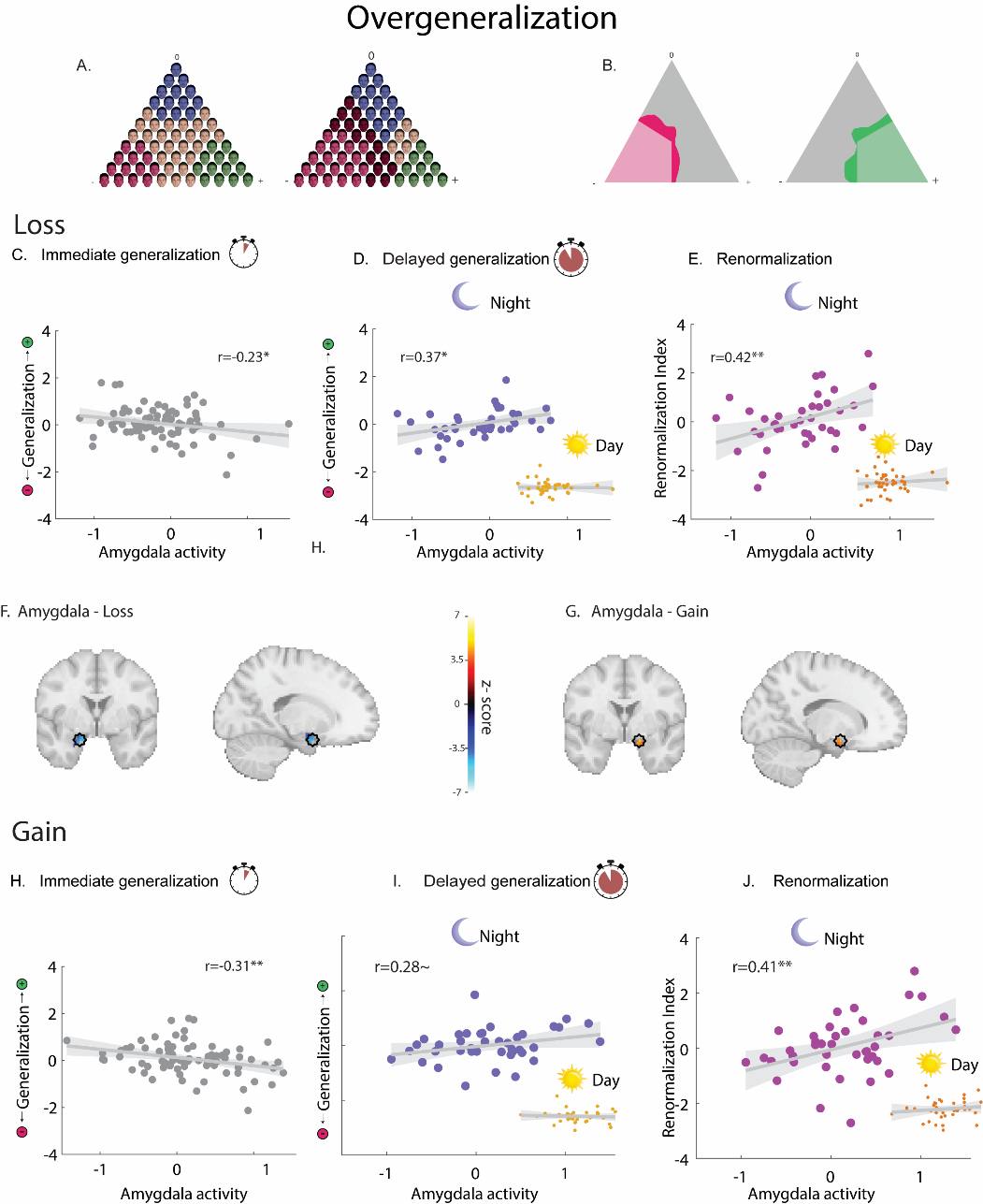

**Supplementary Figure 8. Amygdala activity during learning is associated with immediate overgeneralization and a shift toward positive generalization after sleep. Related to Figure 4.**

Amygdala activity predicts overgeneralization and renormalization. **A–B.** Overgeneralization was defined as responses falling outside correct classification boundaries in the discrimination task, where such errors resulted in point loss. **A.** Left: Schematic of generalization zones surrounding each original face. Right: Example of overgeneralization of the negative face (dark pink=misclassification). **B**. Example participant behavior from two different participants. Light colors indicate correct generalization; dark colors indicate overgeneralization (used for analysis). **C.** Loss: Amygdala activity was negatively correlated with immediate overgeneralization across participants (rₚ=–0.23, p=0.03). **D.** Loss: In the night group, amygdala activity was positively correlated with overgeneralization of the positive face after sleep (rₚ=0.37, p=0.02); no association was observed in the day group. **E.** Loss: In the night group, amygdala activity was positively correlated with the renormalization index, a shift from negative- to positive-face generalization after sleep (rₚ=0.42, p=0.007) ; no association in the day group. **F–G**. Activation maps of amygdala responses to loss (**F**) and gain (**G**) feedback, with black circles representing 3-mm-radius spheres centered on peak voxels used for analysis (for visualization only). **H.** Gain: Amygdala activity was negatively correlated with immediate overgeneralization (rₚ=–0.31, p=0.005). b Gain: In the night group, amygdala activity showed a trend-level positive correlation with renormalization (rₚ=0.28, p=0.07); no association in the day group. **J.** Gain: In the night group, amygdala activity was positively correlated with the renormalization index (rₚ=0.41, p=0.008); no association in the day group.

|  | ***Mean (STD)*** | ***r_p_ Negative (p)*** | ***r_p_ Positive (p)*** | ***r_p_ Neutral (p)*** | ***r_p_ (Generalization score) (p)*** |
| --- | --- | --- | --- | --- | --- |
| ***Time in Bed (min)*** | 437.1 (71.6) | 0.05 (0.77) | -0.04 (0.82) | -0.09 (0.55) | -0.05 (0.78) |
| ***Total Sleep Time (min)*** | 400.2 (69.7) | 0.22 (0.16) | -0.11 (0.48) | -0.09 (0.58) | -0.18 (0.25) |
| ***Sleep Efficiency (%)*** | 92.7 (5.2) | 0.29 (0.06) | -0.13 (0.4) | 0.02 (0.91) | -0.23 (0.14) |
| ***Latency (min)*** | 18.3 (15.5) | -0.29 (0.06) | 0.09 (0.56) | 0.09 (0.56) | 0.2 (0.18) |
| ***N1 (min)*** | 15.1 (11.5) | 0.1 (0.54) | 0.03 (0.87) | -0.19 (0.23) | -0.04 (0.81) |
| ***N2 (min)*** | 150.3 (42.2) | 0.08 (0.64) | 0.05 (0.74) | -0.08 (0.62) | -0.01 (0.94) |
| ***N3 (min)*** | 136.8 (35.2) | -0.11 (0.51) | 0.01 (0.94) | 0.07 (0.67) | 0.06 (0.69) |
| ***REM (min)*** | 96.8 (32.3) | 0.25 (0.11) | -0.24 (0.13) | 0 (1) | -0.27 (0.08) |
| ***Wake (min)*** | 36.8 (24.4) | -0.28 (0.08) | 0.12(0.44) | -0.02 (0.9) | 0.22 (0.17) |
| ***NREM (min)*** | 302.2 (32.3) | 0.018 (0.91) | 0.09 (0.56) | -0.1 (0.54) | 0.04 (0.79) |
| ***WASO (min)*** | 19.7 (17.7) | 0.05 (0.74) | -0.04 (0.82) | -0.06 (0.71) | 0.13 (0.42) |
| ***Number of Arousals (#)*** | 11.1 (5.4) | -0.13 (0.41) | 0.1 (0.52) | -0.13 (0.42) | -0.05 (0.76) |
| ***REM arousals density*** | 0.014 | 0.001 (0.99) | -0.003 (0.98) | -0.045(0.78) | 0.03(0.86) |
| ***Delta (µV^2^/Hz)*** | 115.3 (72.7) | 0.04 (0.83) | -0.13 (0.43) | 0.09 (0.57) | -0.09(0.58) |
| ***Theta (µV^2^/Hz)*** | 5.2 (1.9) | -0.09 (0.59) | -0.04 (0.78) | 0.14 (0.36) | -0.02 (0.89) |
| ***Sigma (µV^2^/Hz)*** | 3.0 (1.5) | -0.13(0.4) | 0.38 (0.01)* | -0.25(0.11) | 0.28(0.07) |

**Supplementary Table 1**. **Sleep architecture and associations with delayed generalization**. **Related to Figure 5 and STAR Methods.**

For each sleep variable, values are reported as the mean and standard deviation (SD) across participants. The table also presents Pearson correlation coefficients (r) and corresponding p-values between each sleep variable and delayed generalization for positive, negative, and neutral faces, as well as the generalization score (positive minus negative face generalization) following sleep. Sleep stages were scored according to standard AASM criteria. Delta-band power (0.75–4 Hz) and sigma-band power (12.5–15 Hz) were quantified during NREM sleep, whereas theta-band power (4–8 Hz) was quantified during REM sleep.
Abbreviations: REM, rapid eye movement; NREM, non-REM sleep; WASO, wake after sleep onset.

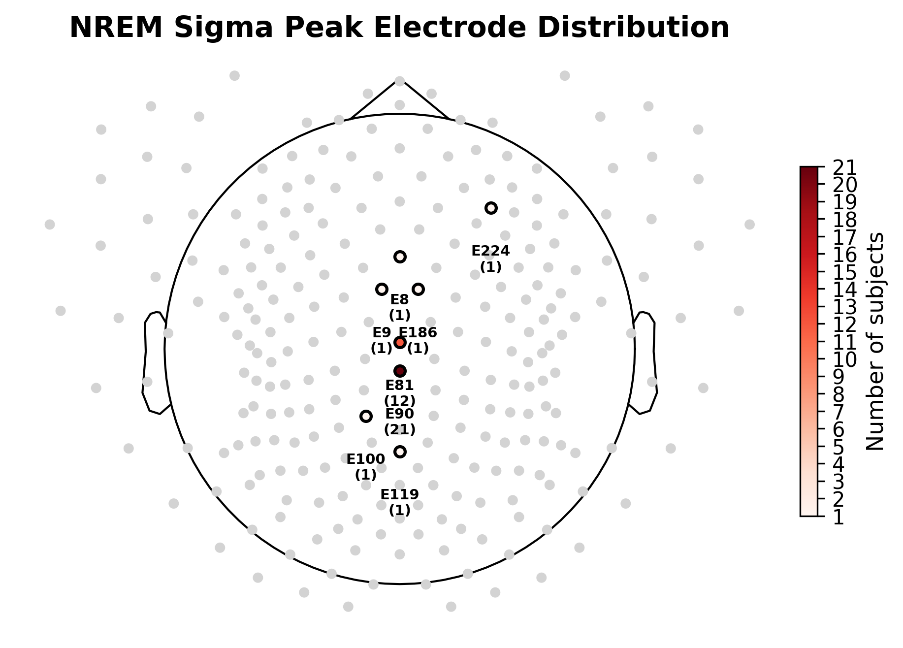

**Supplementary Figure 9. Centroparietal distribution of peak sigma-band activity during NREM sleep. Related to Figure 5 and STAR Methods.**

Histogram depicting the spatial distribution of peak sigma-band electrodes (12–15 Hz, NREM sleep) across participants. The distribution demonstrates a concentration of peak electrodes over centroparietal regions, consistent with the canonical topography of sleep spindles.

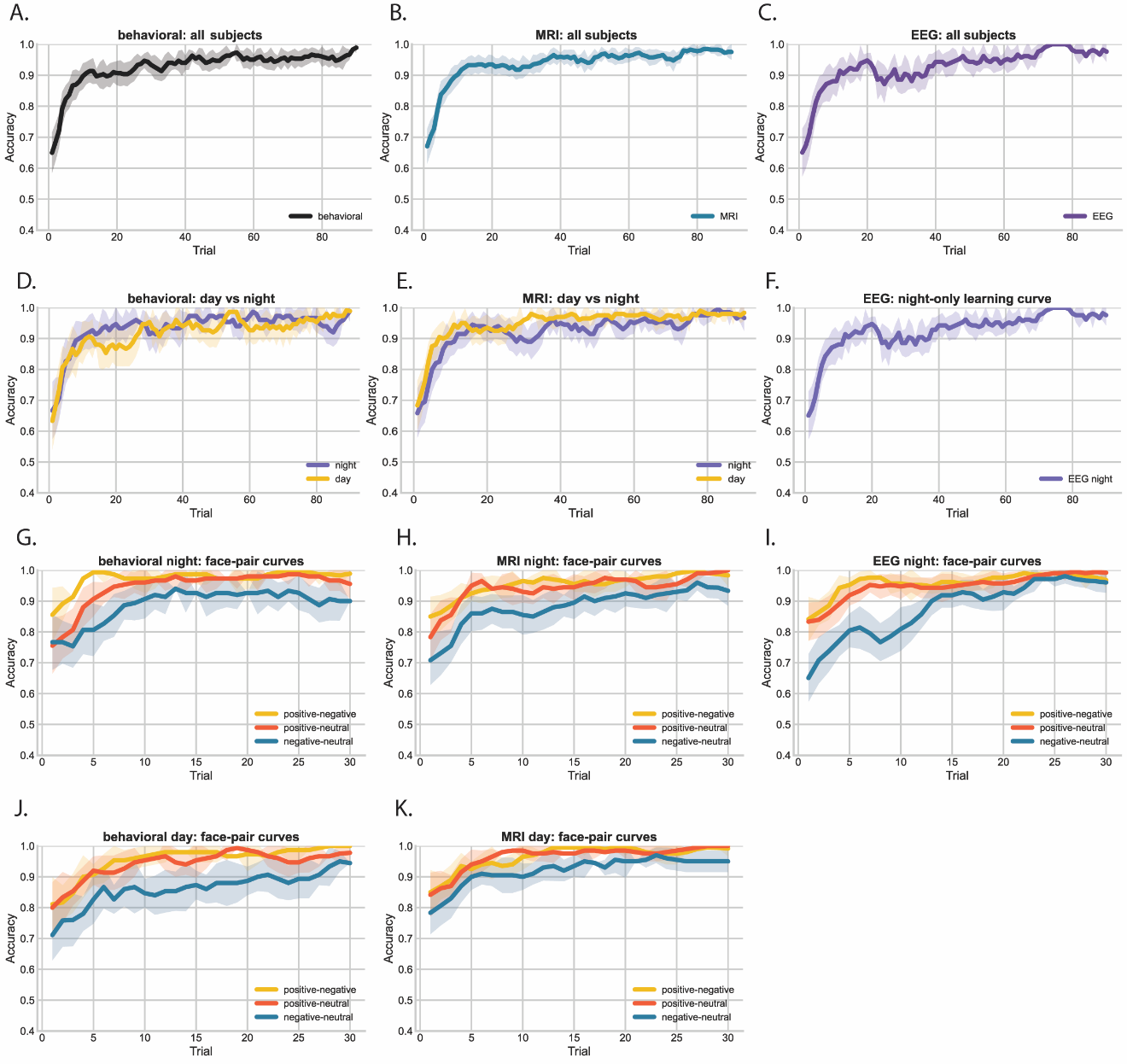

**Supplementary Figure 10. Learning curves across experiments, groups, and face-pair conditions.** **Related to Figure 1.**

Learning accuracy is shown as mean ± SEM across trials, with curves smoothed using a 5-trial moving average. **A–C.** Learning curves for all participants in each experiment: behavioral (**A**), MRI (**B**), and EEG (**C**). **D–F.** Learning curves separated by experimental group (day vs. night) for the behavioral and MRI experiments: behavioral (**D**) and MRI (**E**). The EEG dataset included only a night group (**F**). **G–K.** Learning curves separated by face-pair type. On each learning trial, participants were presented with two faces and were required to choose one of them. Face-pair labels indicate the reinforcement values associated with the two faces presented together: positive-negative (positive and negative faces), positive-neutral (positive and neutral faces), and negative-neutral (negative and neutral faces). Panels **G–I** show the night groups (behavioral **G**, MRI **H**, EEG **I**), and panels **J–K** show the day groups (behavioral **J**, MRI **K**).

Across experiments, participants learned the task successfully. Mean overall accuracy ranged from 91.9% to 94.9% across groups and increased to 95.0%–98.0% during the final 10 trials. Mixed-effects models revealed significant positive learning slopes in the behavioral, MRI, and EEG datasets (all *p* < 0.001). Learning rates did not differ significantly between day and night groups in either the behavioral or MRI experiments (behavioral group × trial: *p* = 0.121; MRI group × trial: *p* = 0.117). Consistent with these findings, summary accuracy comparisons revealed no day–night differences that survived FDR correction. In the MRI experiment, day participants showed numerically higher overall accuracy than night participants (94.9% vs. 92.5%; uncorrected *p* = 0.029, FDR-corrected *p* = 0.293). Learning performance was high across all face-pair types, experiments, and groups. Overall accuracy exceeded 86% for every face pair and reached approximately 91%–99% during the final 10 trials, indicating successful learning even for the most difficult pair type. Within this generally high level of performance, positive-negative and positive-neutral pairs were learned more accurately than negative-neutral pairs. The negative-neutral pair showed the lowest overall accuracy across all groups (behavioral day: 86.3%; behavioral night: 88.8%; MRI day: 92.0%; MRI night: 87.9%; EEG night: 87.5%). Pairwise comparisons showed higher accuracy for positive-negative than negative-neutral pairs in all experiment/group combinations after FDR correction (behavioral night: *p* = 0.0001; behavioral day: *p* = 0.003; MRI night: p = 0.0002; MRI day:p = 0.005; EEG night: p = 0.00007). Positive-neutral accuracy was also higher than negative-neutral accuracy in all groups after FDR correction. Accuracy for positive-negative and positive-neutral pairs was generally similar, except in the behavioral night group, where positive-negative accuracy was modestly higher (*p* = 0.03). Mixed-effects trajectory analyses yielded consistent results, with positive-negative and positive-neutral pairs showing higher accuracy than the negative-neutral reference condition across all experiment/group models.
